# Drought Dominates Engineered Lipid Sink Effects on Sorghum Physiology and Carbon Allocation

**DOI:** 10.64898/2026.05.28.728502

**Authors:** Yuan Liu, Katerina Estera-Molina, Megan Kan, Jordan Masterson, Christina Ramon, Andrew Curtright, Rounak Patra, Lauana Pereira de Oliveira, Kiyoul Park, Edgar B. Cahoon, Wendy H Yang, Jennifer Pett-Ridge

## Abstract

Determining the environmental limits that govern how engineered metabolic traits control whole-plant carbon (C) allocation is essential for developing climate-resilient bioenergy systems. Oil-enhanced sorghum offers a promising strategy to boost aboveground energy density, yet its system-wide effects on root C investment and soil C delivery, particularly under drought, are largely uncharacterized. Here, we provide the first comprehensive assessment of these interactions using whole-plant ^13^CO₂ continuous labeling, depth-specific sampling of roots and rhizosphere soil, and coordinated measurements of photosynthesis, stomatal anatomy, tissue nitrogen, and root non-structural carbohydrates (NSCs). Under well-watered conditions, oil enhancement produced distinct aboveground physiological changes, including slightly longer stomata, lower stomatal density, significantly higher leaf nitrogen concentration and biomass, and greater leaf ^13^C enrichment. Importantly, these aboveground changes did not translate into detectable shifts in the allocation of recent photosynthate to belowground pools. In contrast, drought acted as a dominant regulatory factor, reorganizing C flow across both genotypes by suppressing photosynthesis and leaf water status, increasing root nitrogen and NSC reserves, and significantly promoting the retention of recent assimilates in shallow root systems. Depth-specific ^13^C patterns showed that drought reduced new C incorporation into deep roots while increasing ^13^C enrichment in deep rhizosphere soil, suggesting that drought altered the balance between C investment in root growth and rhizodeposit C inputs at depth. Ultimately, soil moisture availability was an overriding determinant of belowground C partitioning and vertical C delivery, superseding the influence of the engineered lipid sink. These findings provide new mechanistic insight into the environmental constraints on C flow in an engineered bioenergy crop and identify moisture-driven limitations as the primary bottleneck for translating synthetic metabolic innovations into robust, ecosystem-level C outcomes.

## Introduction

Bioenergy crops can play an important role in a sustainable bioeconomy by producing high biomass while contributing to belowground C inputs (Robertson et al., 2017; Jang et al., 2025; Kantola et al., 2025), particularly when grown on marginal lands. Among them, sorghum (*Sorghum bicolor*) is a promising annual feedstock due to its high productivity, high water use efficiency, drought resilience, and deep rooting system (Mullet et al., 2014; Lamb et al., 2022) which can enhance soil C stocks (Lamb et al., 2022). Recent advances in plant metabolic engineering have expanded sorghum’s potential by developing oil-enhanced lines that accumulate triacylglycerols (TAGs) in vegetative tissues, increasing the energy density of aboveground biomass (Park et al., 2025). Field evaluations show that these lines can accumulate up to 5.5% TAG in leaves and 3.5% TAG in stems without large reductions in biomass (Park et al., 2025; Chen et al., 2026). However, establishing a strong lipid C sink may alter whole-plant C and N demand, with potential consequences for how assimilates are allocated among leaves, stems, roots, and rhizosphere soil (Roberts et al., 2025). These effects are important because root and rhizosphere C inputs contribute to mineral-associated organic matter formation (Sokol et al., 2019), a key pathway for long-term soil C persistence (Lavallee et al., 2020). Thus, quantifying how oil enhancement influences whole-plant C partitioning and soil C delivery is essential for evaluating its broader physiological and biogeochemical consequences.

Environmental stresses such as drought interact with plant metabolic traits and are known to dramatically alter plant C allocation patterns (Hasibeder et al., 2015; Wang et al., 2021; Leyrer et al., 2025). Drought typically reduces stomatal conductance, leaf water content, and photosynthetic rate in plants, while increasing intrinsic water-use efficiency and shifting biomass allocation toward roots (Abreha et al., 2022). Reduced sink activity and restricted phloem transport during drought can also lead to the accumulation of non-structural carbohydrates (NSCs) (Qazi et al., 2014) and changes in root N distribution (Cooney et al., 2021), with cascading consequences for belowground C transfer (Hasibeder et al., 2015; Wang et al., 2021). Drought also reshapes rhizosphere inputs by changing the quantity and composition of rhizodeposition (Williams and de Vries, 2020; Ulrich et al., 2022), often reducing total inputs while increasing the proportion of recent assimilates retained belowground (Hasibeder et al., 2015; Brunn et al., 2022). Drought may further increase fine-root turnover (Gaul et al., 2008) and alter the contribution of recent C to soil respiration and microbial uptake (Hagedorn et al., 2016; Hou et al., 2025). Yet, it remains unclear whether oil-enhanced sorghum exhibits altered physiological sensitivity to drought or whether drought modifies the effects of oil enhancement on belowground C routing.

Understanding how recent assimilates move through deep soil profiles is essential for predicting soil C persistence. However, most ^13^CO2 and ^14^CO2 labeling studies focus on surface soils (Ruehr et al., 2009; Hasibeder et al., 2015; Hagedorn et al., 2016; Wang et al., 2021), despite the important role of subsoils in long-term C storage. These studies show that drought can reduce photosynthetic C transfer to soil respiration and microbial biomass while increasing the proportional allocation of recent assimilates to roots and rhizosphere pools (Sanaullah et al., 2012; Burri et al., 2013). Drought also commonly shifts root growth (Kou et al., 2022; Shoaib et al., 2022) and water acquisition toward deeper layers in deep-rooting grasses (Hoekstra et al., 2014; Maan et al., 2023), thereby altering where plant-derived C enters the soil profile and the potential pathways for long-term C stabilization. These depth-dependent processes are especially important in bioenergy sorghum, yet how drought modifies the vertical distribution of newly assimilated C across soil profiles remains critically undercharacterized.

In this study, we used whole-plant ^13^CO₂ continuous labeling, depth-specific sampling of roots and rhizosphere soil, and detailed physiological and biogeochemical measurements to evaluate how the engineered oil sink and drought jointly regulate C allocation in bioenergy sorghum. We hypothesized that: (H1) oil enhancement would increase C assimilation and modify tissue chemistry under well-watered conditions, resulting in greater allocation of recent C to leaves and stems but limited changes in belowground C allocation. (H2) drought would dominate the physiological response, reduce whole-plant C assimilation and shift allocation toward roots, leading to increased root NSCs, root N concentrations, and retention of recent assimilates belowground. (H3) drought would reorganize the vertical distribution of recent C by decreasing deep root allocation while altering the depth at which plant-derived C enters the rhizosphere. To test these hypotheses, we compared the wild-type grain sorghum TX-430 with TZ424-5-3a, an oil-enhanced transgenic line developed in the TX-430 background, under well-watered and drought conditions in a controlled greenhouse experiment. We traced newly fixed C across major organs and soil pools while quantifying physiological traits including photosynthesis, stomatal characteristics, and biomass allocation to assess how these factors shape whole-plant C dynamics.

## 2. Materials and methods

### 2.1 Plant material

This study used two sorghum (*Sorghum bicolor*) genotypes: the wild-type grain sorghum line TX-430 and TZ424-5-3a, an oil-enhanced transgenic line developed in the TX-430 background. TZ424-5-3a was previously engineered to accumulate triacylglycerols in aboveground tissues, thereby increasing the energy density of vegetative biomass (Park et al., 2025). Prior characterization showed that TZ424-5-3a accumulates substantially higher lipid content in leaves and stems than TX-430 without large reductions in biomass, confirming stable expression of the oil-enhancement trait (Park et al., 2025).

### 2.2 Soil sampling and preparation

Soil used in the greenhouse experiment was collected from the Hopland Research and Extension Center of the University of California (39°01.11′N, 123°04.18′W). The site has a Mediterranean climate, characterized by cool, wet winters and hot, dry summers with a mean annual precipitation of 956 mm and mean annual temperature ranging from 7 to 23°C (Foley et al., 2023). The vegetation community is dominated by naturalized annual grasses, including *Avena spp., Aira caryophyllea, Bromus spp., Briza minor* (Bartolome et al., 2007). The soil at the site is classified as a Typic Haploxeralfs of the Witherall-Squawrock complex (Fossum et al., 2022).

In March 2024, after removing surface grass layer (0–3cm), approximately 6,000 lb of soil was excavated using a backhoe tractor from three field soil horizons: 0–20 cm (A), 20–50 cm (B), and 50–100 cm (C). Each horizon was homogenized in the field, placed into sandbags, and transported to the greenhouse facility at the University of California, Berkeley. Soils were air-dried indoors at room temperature, and dried soils at each horizon were combined and thoroughly mixed to produce a uniform composite soil for the greenhouse experiment.

### 2.3 Megacosm preparation, experimental design and sorghum growth

For the greenhouse megacosm experiment, soil profiles were reconstructed to match field bulk density (∼1.5 g cm⁻³). Homogenized soils were packed into 32 clear, impact-resistant polycarbonate cylinders (hereafter referred to as “megacosms”; height 1.22 m, diameter 19.7 cm) (McMaster Carr, Elmhurst, IL). Each megacosm was filled with sequential A, B, C horizons (35 cm, 35 cm, and 40 cm, respectively) packed at field bulk density. Each megacosm contained approximately 56 kg of soil and was sealed at the base with a custom-fitted polycarbonate cap and o-ring (McMaster Carr, Elmhurst, IL).

Soil moisture probes (EC-20; METER GROUP, Pullman, WA) were installed in the A and B horizons to monitor and maintain target soil moisture conditions, while iButton temperature loggers (Maxim Integrated, USA) were embedded at the same depths in half of the megacosms to continuously record soil temperature. Completed megacosms were wrapped in black high-density polyethylene sheeting to prevent algal growth and further enclosed in white polypropylene sacking to reduce heating from solar radiation (Sher et al., 2020). Following assembly, all megacosms were rehydrated to field capacity and equilibrated for two weeks to restore microbial activity and establish uniform soil moisture conditions prior to planting.

The experiment followed a two-factor factorial design with sorghum type (wild-type TX-430 vs. oil-enhanced TZ424-5-3a) and moisture treatment (normal vs. drought) as main factors (n = 8 per treatment; four ¹²C controls and four ^13^C-labeled pots; Fig. 1A). Three seeds were directly sown into each megacosm, covered with a plastic bag to promote germination, and grown under controlled greenhouse conditions (25.6–28.9/20.0–23.3 °C day/night, 16 h photoperiod, ∼50% relative humidity). After germination, seedlings were thinned to one plant per megacosm. All plants were irrigated every other day to maintain near-field moisture capacity for the first 4 weeks. Drought treatment began in week 5 by progressively reducing irrigation to approximately half of that supplied to the well-watered megacosm, imposing a moderate to severe soil water deficit. The well-watered treatments were maintained at around 55–60% of water-holding capacity (WHC), whereas the drought treatment stabilized at approximately 22–30% of WHC.

**Fig. 1.**
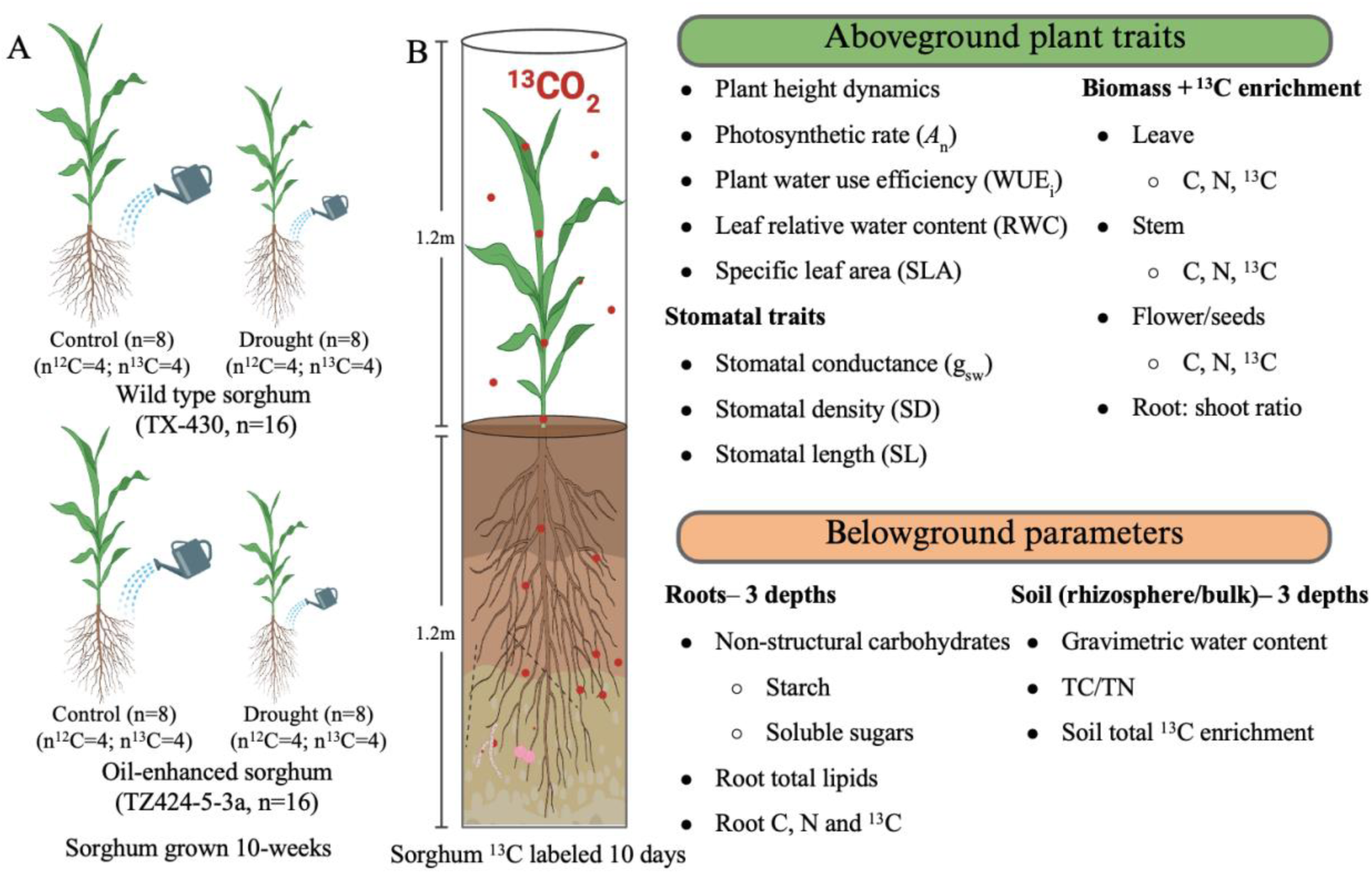
Experimental design and plant–soil parameters measured in a sorghum ¹³CO2-labeling greenhouse experiment. Wild-type (TX-430) and oil-enhanced (TZ424-5-3a) sorghum were grown for 10 weeks under control and soil drought conditions (n = 8 per treatment) with four ¹²CO2 and four ¹³CO2 pots; the latter were pulse-labeled with ¹³CO₂ for 10 days. Aboveground traits included photosynthetic, water-use, and stomatal parameters, while belowground measurements included root non-structural carbohydrates, lipids, C, N, and ¹³C composition, and soil physicochemical and microbial properties across three depths. TC=total carbon, TN=total nitrogen.

Plant height was measured weekly from the soil surface to the tallest leaf. Visual growth and canopy characteristics were recorded to track physiological responses throughout the 10-week growth period.

### 2.4 ^13^CO₂ continuous -labeling experiment

After 10 weeks of growth, a 10-day ^13^CO₂ continuous labeling was performed to trace the assimilation and allocation of newly fixed carbon within plant and soil pools (Fig. 1). Each megacosm was enclosed in a transparent impact-resistant polycarbonate cylinder (height 1.22 m, diameter 19.7 cm) (McMaster Carr, Elmhurst, IL) fitted to the pot. Half of the megacosms from each treatment were labeled with natural abundance ^12^CO2 (Praxair, Danbury, CT) as controls, while the other half received 98 atom % ^13^CO₂ (Sigma-Aldrich). The chambers were connected to an automated CO2 supply and monitoring system, maintaining average CO2 concentrations between 642–1062 ppm during the light period.

Labeling was conducted under greenhouse light conditions (∼700 µmol m⁻² s⁻¹ PAR) for 16 h per day. ^13^CO₂ concentrations and isotopic composition within the ^13^C labeled chambers were continuously monitored using a Picarro G2201-i Analyzer (Picarro Inc., Santa Clara, CA, USA), and CO2 concentrations within the ^12^C labeled chambers were continuously monitored using a SBA-5 infrared gas analyzer (SBA-5; PP Systems, Amesbury, MA). Both CO2 analyzers interfaced with a CR1000 datalogger (Campbell Scientific Logan, UT, USA) used both for data accumulation and CO2 control. This setup enabled real-time tracking of ¹²CO₂ and ^13^CO₂ concentrations across all 32 chambers (16 labeled and 16 controls). Detailed description of the labeling system and operation is provided in previous studies using the same setup (Sher et al., 2020; Sokol et al., 2024)

### 2.5 Plant physiological and morphological measurements

#### 2.5.1 Leaf gas exchange measurement

One week prior to ^13^C labeling, leaf gas exchange parameters, including steady-state photosynthetic rate (*A*ₙ) and stomatal conductance (*g*sw), were measured on the third youngest fully expanded leaf of each plant using a LI-6800 Portable Photosynthetic System equipped with a standard 6 cm^2^ cuvette (LI-COR Inc., Lincoln, NE). Measurements were conducted between 10:00-14:00 under the controlled chamber conditions: leaf temperature 27°C, relative humidity 65%, photosynthetic photon flux density (PPFD) 1800 µmol m^-2^ s^-1^, reference CO₂ concentration 420 µmol mol^-1^, and flow rate 500 µmol s^-1^. Intrinsic water-use efficiency (WUEᵢ) was calculated as the ratio of *A*ₙ to *g*ₛw.

#### 2.5.2 Leaf relative water content (RWC) and specific leaf area (SLA)

At harvest, 12 leaves per plant were cut into approximately 1 × 1.5-inch sections and fresh weight was recorded immediately. Samples were then placed into 50-ml centrifuge tubes filled with deionized water until the leaf pieces were fully submerged. After 4–6 h of hydration, the water was gently poured off, and leaves were carefully dried using paper towels to remove surface moisture before measuring saturated weight. Samples were then oven-dried at 70 °C for 24 h to obtain dry weight. RWC was calculated as:

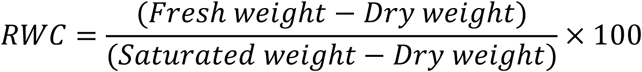

SLA was determined as the ratio of leaf area to leaf dry weight (cm^2^ g^-1^) as measured above.

#### 2.5.3 Stomata traits measurements

Stomatal density and length were quantified on both the adaxial and abaxial surfaces of the third youngest fully expanded leaf from each sampled plant using clear nail-polish imprints and light microscopy. At harvest, transparent nail polish was applied to the middle section of the third youngest fully expanded leaf on both upper and the lower surfaces. After approximately 1 min, transparent adhesive tape was gently pressed onto the dried polish to ensure full contact, then peeled off and mounted onto a glass microscope slide.

Stomatal density was then determined under a stereomicroscope (×400 magnification; Zeiss^®^ Axio Zoom V16, Jena, Germany) by counting the number of stomata within a defined area of 883 × 648 µm. Three non-overlapping fields of view were analyzed per sample, and the mean value was used for each replicate. Stomata length was measured from the same images using ImageJ software (version 1.54g, Java 1.8.0_345, Wayen Rasband, US National Institutes of Health, Bethesda, MD, USA). Average stomatal density and stomatal length were calculated as the mean of adaxial and abaxial measurements for each leaf.

### 2.6 Plant and soil sampling and elemental and isotope measurements

At harvest, all plants were clipped at the base of the stems, and separated into leaves, stems, and flowers, when present. Roots were further divided by depth intervals (0–35 cm, 35–70 cm, and 70–110 cm) and all visual coarse roots were carefully picked out. A subset of roots was flash frozen in liquid nitrogen and stored at –80°C for subsequent total lipids analysis. For each soil horizon, roots were washed in deionized water and dried with paper towels. All plant materials were oven-dried at 65 °C to constant weight, and dry mass was recorded. A representative subset of each tissue was then ground to a fine powder using a ball mill (Retsch MM400 mixer mill, Germany) and stored in airtight scintillation vials containers for elemental (carbon (C), nitrogen (N)) and isotopic analysis.

Rhizosphere (< 2 mm from the root) and bulk soil (> 2mm from the root) were collected from each depth. Rhizosphere soil was collected by gently shaking roots to detach soil adhering to the root surface, whereas bulk soil was collected from adjacent regions without visible root contact. A subset of rhizosphere and bulk soil samples was immediately transferred into sterile polyethylene bags, flash-frozen on dry ice and stored at −80 °C until further analysis. A subset of 10 g of soil was used to determine gravimetric water content after drying at 105 °C for 24 h, and the remaining soil was finely ground for elemental (C, N) and isotopic analysis.

C and N concentrations of plant and soil samples were measured using an Elementar vario EL Cube CHNS analyzer (Elementar, Germany) with sulfanilamide as standard. Stable carbon isotope composition (δ^13^C) was determined using an elemental analyzer coupled to an isotope ratio mass spectrometer (EA–IRMS; Costech ECS 4010, Costech Analytical Technologies, Valencia, CA, USA) at the Stable Isotope facility, Yale University. Atom% ^13^C enrichment was calculated relative to unlabeled controls, and the amount of newly assimilated 13C in each fraction was determined by multiplying atom% excess by total C content. Total ^13^C retention was expressed both as absolute incorporation (mg ^13^C) within individual pools and as fractional allocation (%) among plant tissues and soil compartments.

### 2.7 Rhizodeposition index calculation

The rhizodeposition index (RDI) was calculated to quantify the proportion of recent assimilates retained in soil relative to roots at each depth. For each depth, we summed excess ¹³C recovered in rhizosphere and bulk soil and in roots, and calculated:

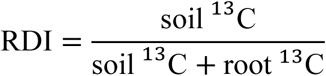

where soil ^13^C represents the total excess ^13^C recovered in rhizosphere and bulk soil fractions, and root ^13^C represents excess ^13^C recovered in root tissue. This index reflects the fraction of retained, non-respired recent assimilates present in soil relative to roots within each depth. Higher RDI values indicate a greater proportion of retained recent assimilates in soil relative to roots at a given depth.

### 2.8 Non-structural carbohydrates

Root non-structural carbohydrates (NSC) were quantified as soluble sugars (glucose, fructose, and sucrose) and starch using freeze-dried root material from each harvested plant.

Freeze-dried roots were finely ground, soluble sugars were extracted and quantified from the supernatant, and the remaining insoluble pellet (after soluble sugar removal) was used for enzymatic starch determination. Total soluble sugars were calculated as the sum of glucose, fructose, and sucrose, and total NSC was calculated as total soluble sugars plus starch. Starch extraction and quantification followed Amaral et al. (2007) with modifications as described below.

#### 2.8.1 Soluble sugar content (glucose, fructose, and sucrose)

Soluble sugars (glucose, fructose, and sucrose) were extracted from freeze-dried root tissue using hot ethanol. Samples were incubated in 80% (v/v) ethanol buffered with HEPES (pH 7.8) at 80°C for 20 min, and the supernatant was collected. The extraction was repeated twice with 80% buffered ethanol and once with 50% HEPES-buffered ethanol. Supernatants were pooled, and the total extraction volume was recorded. Sugar concentrations were quantified using an enzymatic assay coupled to NADP+ reduction and measured in a microplate reader at 340 nm. The assay buffer contained 0.1 M HEPES (pH 7.0), MgCl₂, ATP, NADP+, and glucose-6-phosphate dehydrogenase (G6PDH). Glucose was quantified following addition of hexokinase; fructose was quantified following addition of phosphoglucose isomerase (PGI), and sucrose was quantified after hydrolysis with invertase. At each step, formation of NADPH was monitored as the increase in absorbance at 340 nm. Concentrations were calculated from glucose standard curves (0.01–5 mM) and corrected for extraction volume and dilution factors.

#### 2.8.2 Starch content

After removal of soluble sugars, pellets were treated with 120 U mL⁻¹ of α-amylase (EC 3.2.1.1; Megazyme®, Australia) diluted in 10 mM MOPS buffer (pH 6.5) at 75 °C for 1 h, followed by the addition of 30 U mL⁻¹ of amyloglucosidase (EC 3.2.1.3; Megazyme®, Australia) diluted in 100 mM sodium acetate buffer (pH 4.5) at 50 °C for 1 h. The glucose released from starch hydrolysis was quantified using the Megazyme D-Glucose Assay Kit (K-GLUC) at 45 °C for approximately 20 min, and absorbance was measured at 510 nm using glucose (1 mg mL⁻¹) as a standard. Starch concentrations were expressed on a dry mass basis (freeze-dried root mass).

### 2.9 Root total lipids measurement

Root total lipids were extracted and quantified for each plant following the modified Bligh–Dyer method as described in Park et al. (2025), using triheptadecanoin as an internal standard and gas chromatographic analysis of fatty acid methyl esters (FAMEs) for quantification.

### 2.10 Statistical analysis

All statistical analyses were performed in R (version 4.4.0, R Core Team, 2024). We used two-way analysis of variance (ANOVA) to test the main and interactive effects of sorghum type (wild-type vs. oil-enhanced) and moisture treatment (well-watered vs. drought) on plant and soil response variables. Prior to analysis, data were examined for normality (Shapiro–Wilk test) and homogeneity of variances (Levene’s test); variables that violated assumptions were log-transformed as needed. When a significant interaction between sorghum type and moisture treatment was detected (*p* < 0.05), simple-effects tests were performed using estimated marginal means (*emmeans*) to compare sorghum types within each moisture treatment (Tukey-adjusted pairwise contrasts). When the interaction was not significant, main effects were interpreted based on the two-way ANOVA, and pairwise comparisons of marginal means were conducted to identify significant differences between factor levels. Differences were considered statistically significant at *p* < 0.05 and considered as marginal when 0.05 ≤ *p* < 0.10. In figures, significance is indicated as follows: *p* < 0.05 (*), 0.05 ≤ *p* < 0.10 (†), and *p* > 0.10 (ns). All summary statistics and plots show mean ± standard error (SE).

Rhizodeposition index (RDI) was analyzed using a linear mixed model with moisture treatment, sorghum type, and depth as fixed effects and pot as a random effect. Given the strong interaction between moisture treatment and depth, depth effects were evaluated within each moisture level using estimated marginal means with Tukey-adjusted contrasts, and moisture effects were similarly tested within each depth. Pearson correlation coefficients were used to examine relationships among physiological traits and ^13^C allocation patterns across plant and soil fractions, and the resulting correlation matrix was visualized using the *corrplot* package in R.

## 3. Results

### 3.1 Soil moisture, plant height dynamics and root lipids

Drought reduced soil gravimetric water content (GWC) by approximately 50% across all depths, confirming successful moisture treatment separation (Figs. S1–S2). Surface soils (0–35 cm) exhibited greater temporal variability than deeper layers, while oil enhancement did not influence soil moisture dynamics (Figs. S1–S2). Plant height followed a sigmoidal growth pattern and diverged between treatments after ∼50 days, with sorghum plants grown under drought conditions reaching significantly lower final heights in both genotypes (Fig. S3).

Oil enhancement substantially increased total root lipid content under both moisture conditions (Fig. S4A). Under well-watered conditions, oil-enhanced sorghum accumulated more than twice the total lipid content of wild-type plants (5.51 *vs*. 2.72 µg mg⁻¹ dw). This pattern persisted under drought, with oil-enhanced sorghum roots containing ∼45% more total lipids than wild-type roots (5.51 *vs*. 3.04 µg mg⁻¹ dw). Fatty acid composition differed consistently between sorghum genotypes across both moisture conditions. Oil-enhanced sorghum exhibited a higher proportional abundance of C18:1, while showing lower proportions of the remaining fatty acids (C18:3, C18:2, C18:0, and C16:0) compared to wild-type plants under both well-watered and drought conditions (Fig. S4B). Overall, these results demonstrate that oil enhancement markedly elevates root total lipids and shifts fatty acid distribution, while genotype-driven compositional patterns remain consistent across moisture treatments.

### 3.2 Plant physiological and morphological responses to oil enhancement and drought

Drought significantly reduced leaf relative water content (RWC), steady-state photosynthetic rate (*A*ₙ), and stomatal conductance (*g*sw), while significantly increasing specific leaf area (SLA), root non-structural carbohydrates (NSC), and intrinsic water-use efficiency (WUEᵢ) (Fig. 2). Oil enhancement did not significantly alter any of these parameters under either moisture regime. In addition to increasing total root NSC concentrations, drought significantly increased root glucose, fructose, sucrose, and starch concentrations (Fig. S5). Sucrose was the dominant soluble sugar and accounted for most of the drought-associated NSC accumulation, whereas glucose and fructose comprised a comparatively small fraction of the soluble sugar pool. Starch also increased under drought but remained low relative to soluble sugars.

**Fig. 2.**
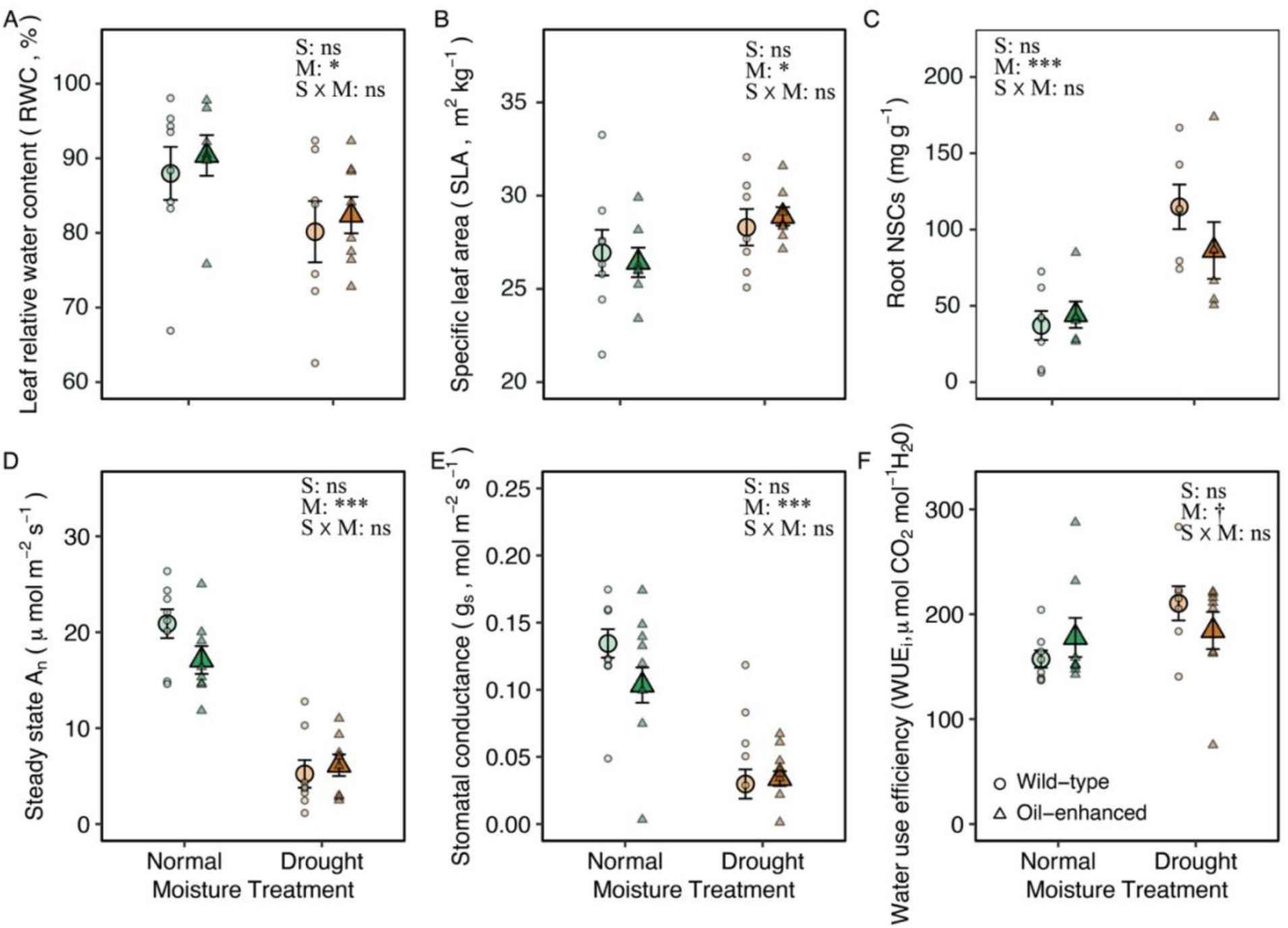
Effects of oil enhancement and drought on sorghum plant physiology parameters. Enlarged circle and triangles show mean ± SE (n = 8). Circles and triangles represent wild-type (TX-430) and oil-enhanced (TZ424-5-3a), respectively. Green and brown colors denote normal and drought soil moisture treatments. Results of two-way ANOVA indicate effects of sorghum type (S), moisture (M), and their interaction (S × M). Symbols: * *p* < 0.05; † *p* < 0.1; ns = not significant.

Stomatal length and density were primarily influenced by moisture treatment (Fig. 3). Drought significantly reduced stomatal length on both the adaxial and abaxial surfaces and marginally increased stomata density. In contrast, oil-enhanced sorghum showed the opposite tendency, producing longer and lower stomatal density than wild-type sorghum plants, particularly under well-watered conditions. On average, oil-enhanced sorghum had 5% longer and 20% less dense stomata than wild-type sorghums under well-watered conditions (Fig. C, F).

**Fig. 3.**
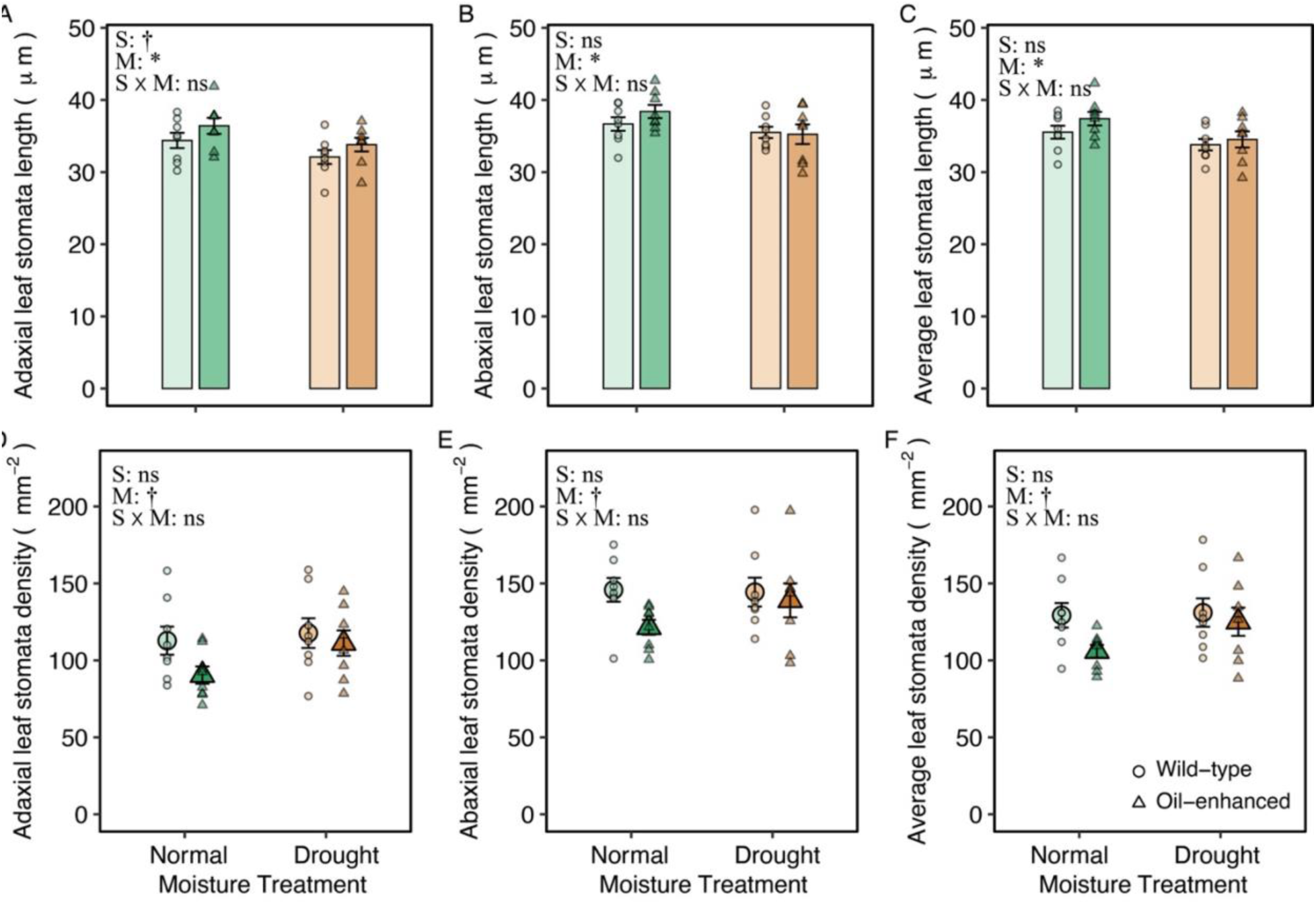
Effects of oil enhancement and drought on sorghum leaf stomatal traits. (A–C) Stomatal length on the adaxial surface (A), abaxial surface (B), and averaged across both surfaces (C). (D–F) Stomatal density on the adaxial surface (D), abaxial surface (E), and averaged across both surfaces (F). Bar and enlarged point show mean ± SE (n = 8). Circles and triangles represent wild-type and oil-enhanced sorghum, respectively. Green and brown colors denote normal and drought moisture treatments. Results of two-way ANOVA indicate effects of sorghum type (S), moisture (M), and their interaction (S × M). Symbols: * *p* < 0.05; † *p* < 0.1; ns = not significant.

### 3.3 Plant elemental composition and biomass allocation under oil enhancement and drought

Plant tissue chemistry responded differently to drought and oil enhancement treatment (Fig. 4). Drought significantly increased N concentrations in all tissues, especially stems and roots, while having minimal effects on C concentration; consequently, C:N ratios decreased. Oil enhancement significantly increased leaf N concentration by 38% (Fig. 4D) and lowered C:N ratios (27%) only under well-watered conditions (Fig. 4G). Consistent with these patterns, drought significantly reduced plant height, leaf, stem, shoot, and root biomass, while increasing the root-to-shoot ratio (Fig. 5). Oil enhancement significantly enhanced leaf biomass by 26% under well-watered conditions (Fig. 5B) but had no effect under drought.

**Fig. 4.**
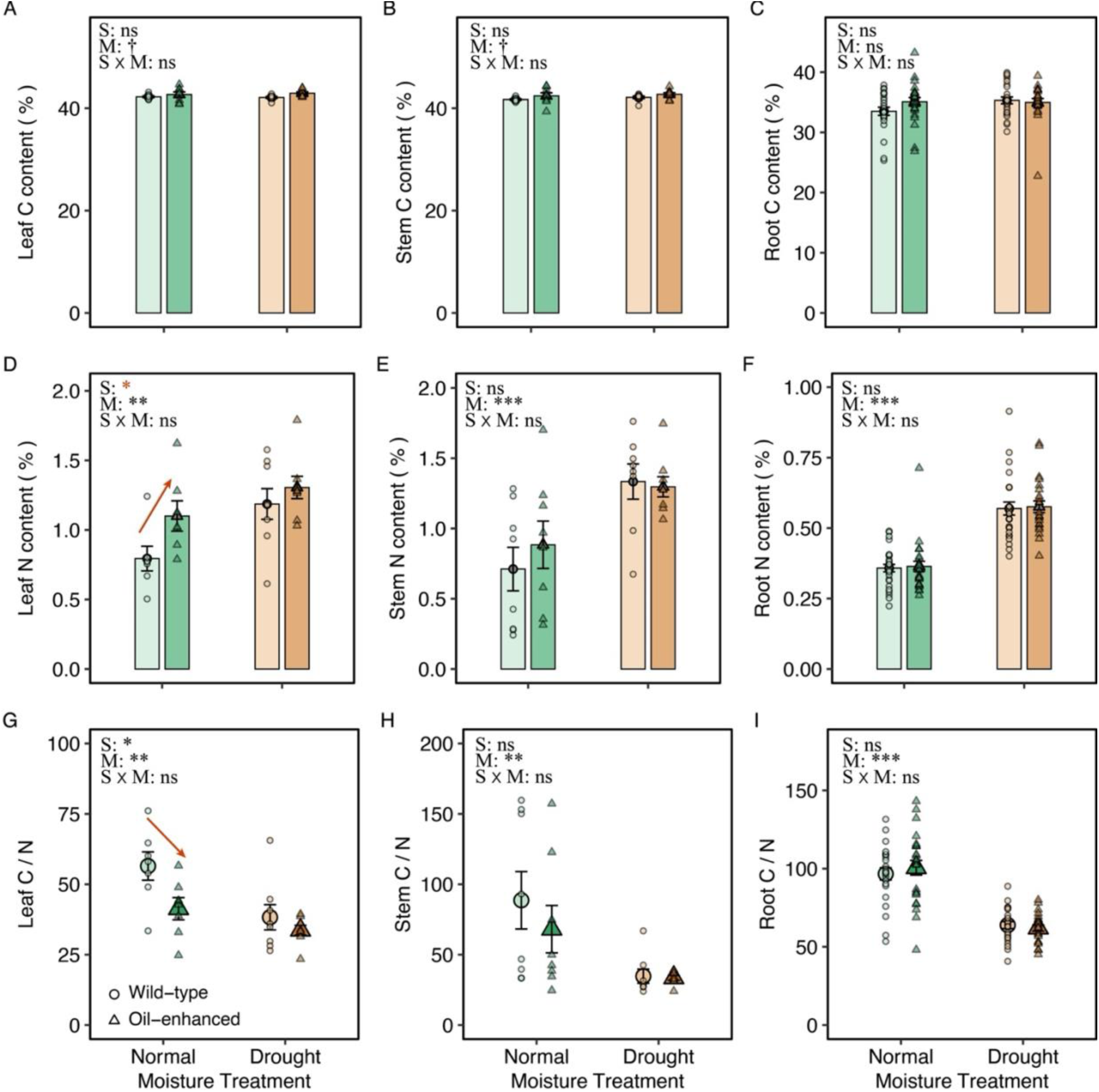
Effects of oil enhancement and drought on sorghum tissue carbon and nitrogen concentrations. (A–C) Leaf, stem, and root C content (%). (D–F) Leaf, stem, and root N content (%). (G–I) Leaf, stem, and root C:N ratios. Bars and enlarged points show mean ± SE (n = 8 for leaf and stem, n=24 for root across all depths). Circles and triangles represent wild-type and oil-enhanced sorghum, respectively, under control (green) and drought (brown) conditions. Results of two-way ANOVA indicate effects of sorghum type (S), moisture (M), and their interaction (S × M). Symbols: ** *p* < 0.01; * *p* < 0.05; † *p* < 0.1; ns = not significant.

**Fig. 5.**
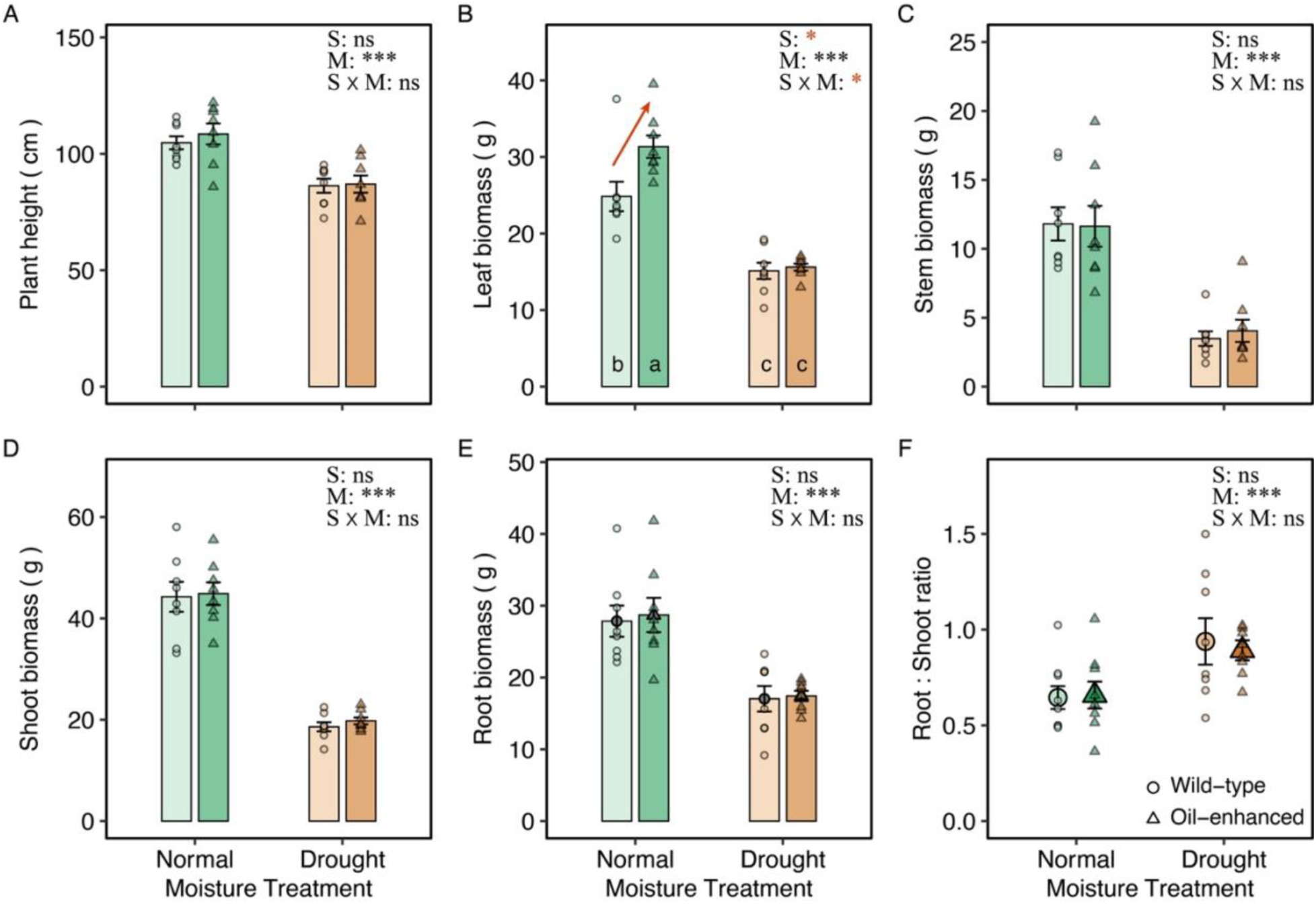
Effects of oil enhancement and drought on sorghum growth and biomass allocation. (A) Plant height, (B) leaf biomass, (C) stem biomass, (D) total shoot biomass, (E) root biomass, and (F) root-to-shoot ratio of wild-type (circles) and oil-enhanced (triangles) sorghum grown under normal and drought moisture treatments. Bars and enlarged points represent means ± SE (n = 6–8). Green and brown colors denote normal and drought treatments, respectively. Results from two-way ANOVA show effects of sorghum type (S), moisture (M), and their interaction (S × M). Symbols: * *p* < 0.05; ** *p* < 0.01; *** *p* < 0.001; ns = not significant.

### 3.4 Plant ^13^C distribution and new photosynthate allocation under oil enhancement and drought

Drought significantly increased ^13^C enrichment in leaves and topsoil roots but reduced 13C enrichment in deeper roots, while simultaneously enhancing ^13^C enrichment in deep rhizosphere soil (Fig. 6). Under well-watered conditions, oil enhancement modestly but consistently increased leaf ^13^C enrichment, by approximately 63% relative to wild type, although sorghum genotype did not exert a significant overall effect on ^13^C enrichment in any plant tissue or soil. Root ^13^C enrichment showed contrasting depth patterns across moisture treatments. Under drought conditions, ^13^C enrichment was highest in the topsoil and decreased with depth, whereas under well-watered conditions it was lowest in the topsoil and increased with depth. Stem ^13^C enrichment did not differ significantly between moisture treatments or genotypes.

**Fig. 6.**
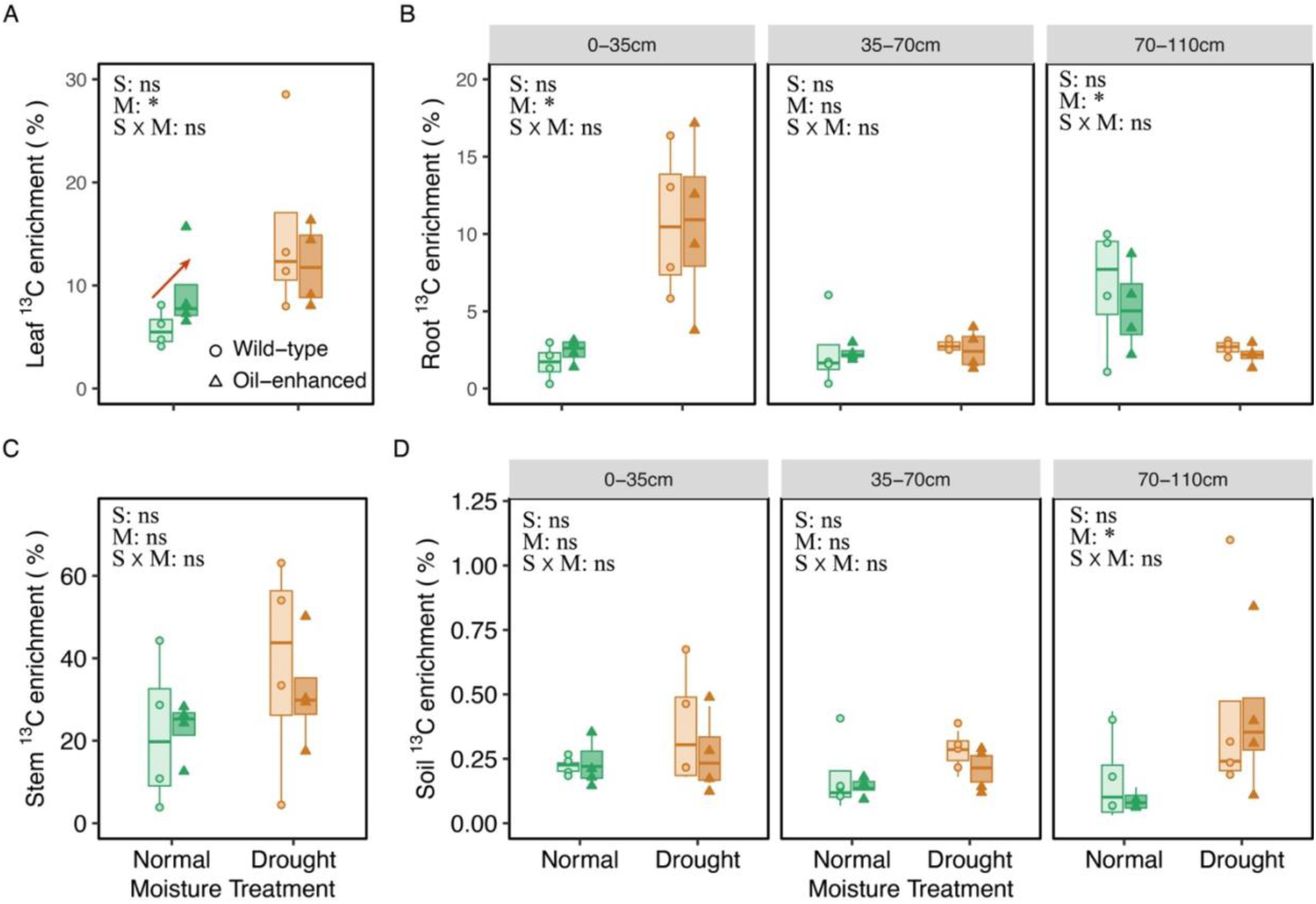
Effects of oil enhancement and drought on ¹³C enrichment in sorghum tissues and rhizosphere soil. (A) Leaf, (B) root, (C) stem, and (D) rhizosphere soil ¹³C enrichment (%) of wild-type (circles) and oil-enhanced (triangles) sorghum grown under normal and drought conditions in a CA annual grassland soil (Typic Haploxeralf). Soil and root ¹³C enrichment were measured across three depth intervals (0–35, 35–70, and 70–110 cm). Green and brown colors represent normal and drought moisture treatments, respectively. Boxes indicate interquartile ranges with median lines, and whiskers show data range. Two-way ANOVA results indicate effects of sorghum type (S), moisture (M), and their interaction (S × M). Symbols: * *p* < 0.05; ns = not significant.

Total ^13^C retention was reduced by 14% and 43% by drought for wild-type and oil-enhanced sorghum, respectively. Under well-watered conditions, oil-enhanced sorghum assimilated 23% more ^13^C than wild-type (Fig. 7A). Most ^13^C was recovered in leaves and stems, whereas roots and soil accounted for smaller proportions. Proportionally, a significantly greater fraction of ^13^C was allocated to roots under drought in both sorghum types (Fig. 7B). In contrast, neither sorghum type nor moisture treatment significantly affected ^13^C retention in soil. On average, approximately 18% of total assimilated ^13^C was recovered in soil, independent of genotype or treatment.

**Fig. 7.**
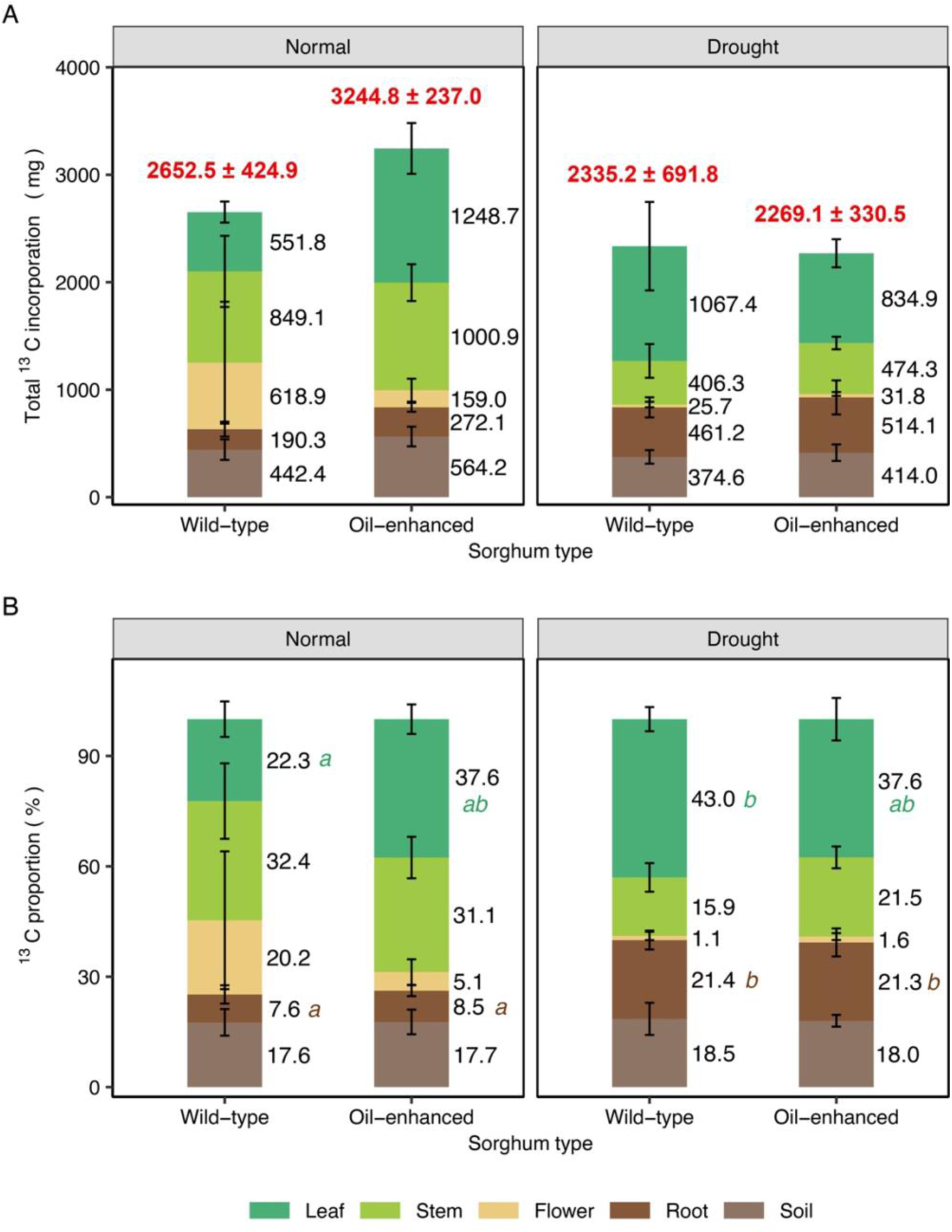
Effects of oil enhancement and drought on total and proportional ¹³C incorporation into sorghum tissues and soil. (A) Total ¹³C incorporation (mg) and (B) proportional ¹³C distribution (%) across plant tissues (leaf, stem, flower, root) and soil for wild-type and oil-enhanced sorghum under normal and drought moisture treatments. Bars represent means ± SE (n = 4). Numbers within each stacked bar denote the mean ¹³C content (A) or percentage contribution (B) of each component. Red numbers on top of each bar indicate total ¹³C incorporation (mean ± SE). Different lowercase letters denote significant differences among treatments within each fraction based on two-way ANOVA followed by *post hoc* tests (*p* < 0.05).

The rhizodeposition index (RDI) reflects the fraction of retained, non-respired recent assimilates present in soil relative to roots within each depth. RDI varied strongly with both depth and moisture treatment (Fig. 8). Under well-watered conditions, RDI declined modestly with increasing depth, whereas under drought it increased significantly across the soil profile. Drought resulted in substantially lower rhizodeposition in surface soil but higher values at depth. Sorghum genotype had limited influence and differed only at the mid-depth under drought, where oil-enhanced sorghums showed a slightly elevated RDI.

**Fig. 8.**
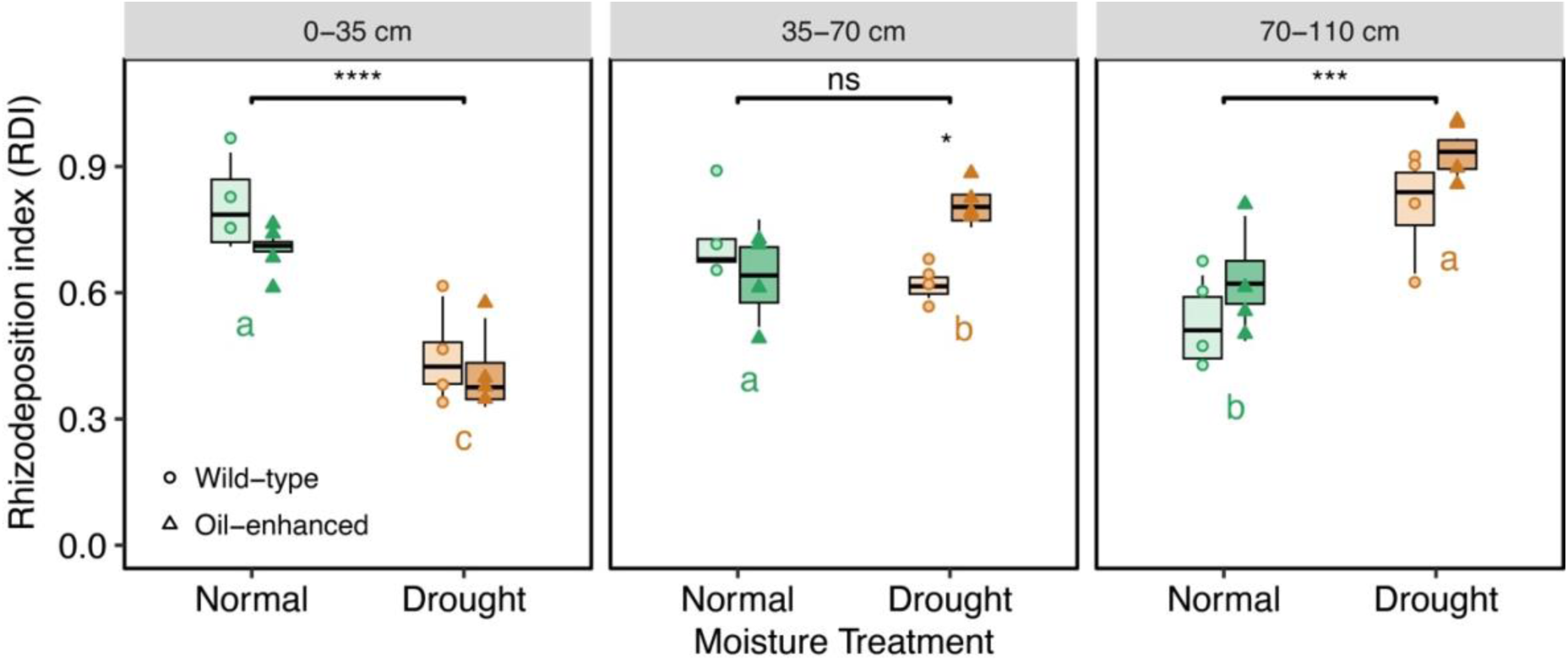
Rhizodeposition index (RDI, soil ¹³C / [soil ¹³C + root ¹³C]) for wild-type and oil-enhanced sorghum across soil depths under normal and drought moisture conditions. Each point represents an individual replicate value of each depth and treatment. Boxes indicate interquartile ranges with median lines, and whiskers show data range. Horizontal brackets indicate moisture effects within a given depth, and letters show Tukey-adjusted depth differences within each moisture treatment based on estimated marginal means from the mixed model (*p* < 0.05); Significance levels: **** *p* < 0.0001, *** *p* < 0.001, * *p* < 0.05; ns = not significant.

Root non-structural carbohydrate concentration was strongly correlated with root N (R² = 0.84, p < 0.01; Fig. S6), indicating tight coupling between nitrogen status and carbohydrate storage across treatments.

### 3.5 Correlations between new photosynthate allocation and physiological traits

Correlation analysis revealed distinct physiological controls on the allocation of new ^13^C among plant and soil pools (Fig. S7). The fraction of new ^13^C in leaves (fLeaf) was positively correlated with stomatal density and root N but negatively correlated with stomatal length, leaf RWC, and soil GWC. The fraction in roots (fRoots) was positively correlated with root-to-shoot ratio, root N, and adaxial stomatal density, and negatively with GasEx, gₛ, leaf RWC, and soil GWC. The fractions in stems and flowers (fStem, fFlower) were correlated only with specific leaf area (SLA), negatively for stems and positively for flowers, while the fraction in soil (fSoil) was negatively correlated with leaf N and SLA.

## 4. Discussion

### 4.1 Effects of Oil Enhancement on Plant Physiology and Plant Carbon Allocation

Oil enhancement altered sorghum physiological traits in a distinctly moisture-dependent manner (Fig. 9). Under well-watered conditions, oil-enhanced sorghum exhibited slightly longer stomata, reduced stomatal density, significantly higher leaf N concentration, and greater leaf biomass compared with the wild-type. These coordinated shifts were supported by isotopic evidence, as oil-enhanced sorghum had greater ^13^C enrichment in leaves, indicating enhanced assimilation and retention of recent photosynthates when water was not limiting. Together, these results confirm our first hypothesis (H1), showing that increased vegetative oil accumulation was associated with altered aboveground C and N allocation under well-watered conditions, when soil moisture availability supports high metabolic activity. In contrast, drought imposed overriding constraints on a full range of plant physiological processes, including photosynthesis, stomatal regulation, biomass production, and leaf water status, suppressing genotypic differences and underscoring that soil moisture is a primary regulator of physiological trait expression in sorghum.

**Fig. 9.**
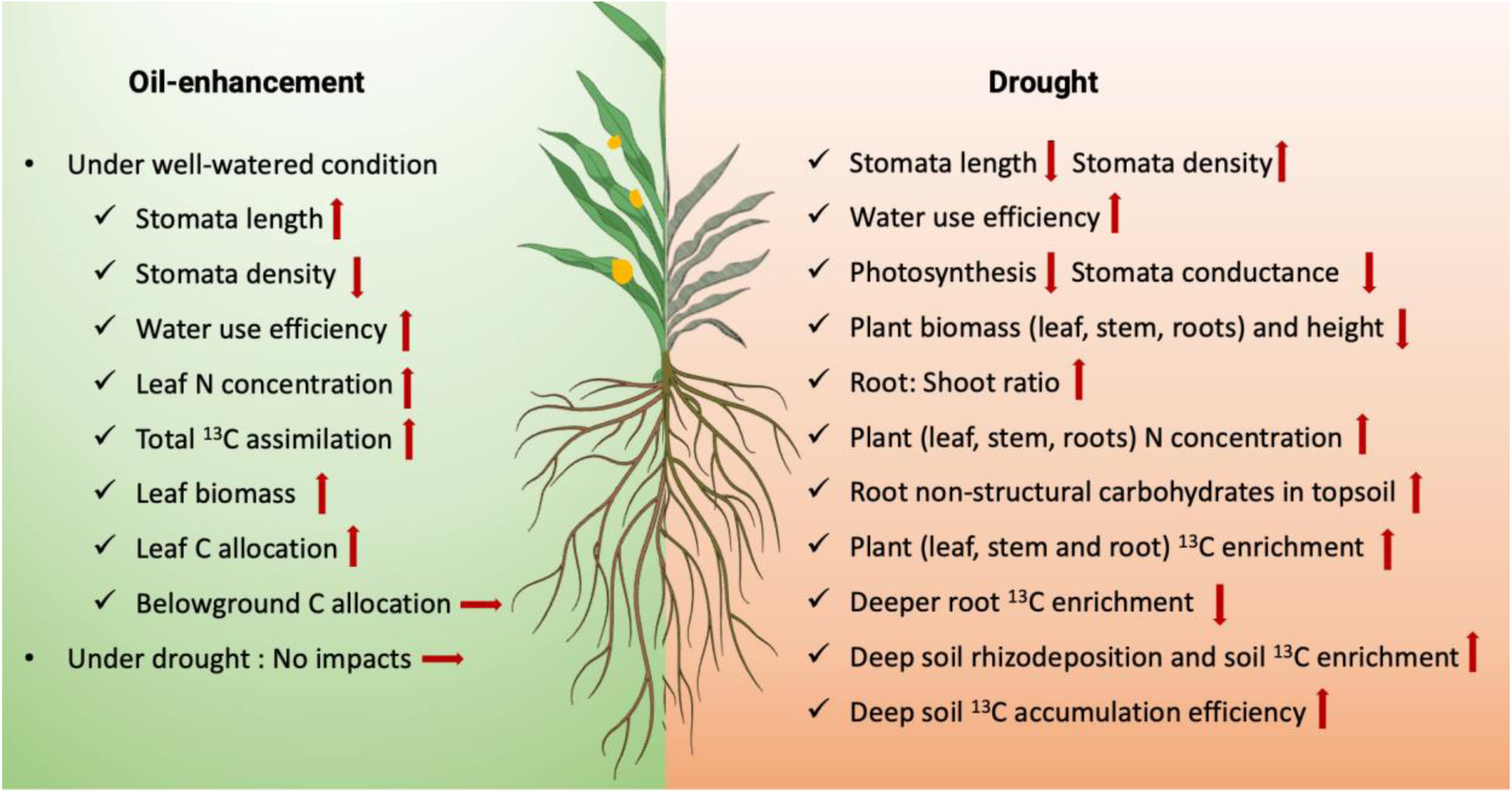
Conceptual summary of oil enhancement and drought effects on sorghum physiology and carbon allocation. Schematic illustration summarizing the effects of oil enhancement (left, green) and drought (right, orange) on sorghum stomatal traits, physiology, nitrogen status, and carbon allocation. Arrows indicate the direction of change (↑ increase, ↓ decrease, → no change). Red asterisks indicate statistically significant effects at *p* < 0.05; on the oil-enhancement side, asterisks refer to significant genotype effects under well-watered conditions, whereas on the drought side, asterisks refer to significant moisture effects.

Leaf N is essential for sustaining photosynthetic efficiency and providing the energetic cofactors required for fatty-acid and triacylglycerol production (Walker et al., 2020). In our study, we measured significantly higher leaf N concentration in oil-enhanced sorghum under well-watered conditions, which likely reflects the increased N demand required to support elevated lipid biosynthesis. This interpretation is supported by recent multisite field evidence showing that TX430-derived oil sorghum lines had higher tissue N concentrations than wild-type plants, suggesting that increased N demand may be a consistent consequence of vegetative lipid accumulation in this genetic background (Chen et al., 2026). This is also consistent with previous evidence that high-lipid crops have enhanced N acquisition and allocate proportionally more N to vegetative tissues to sustain photosynthesis and lipid-producing metabolic pathways

(Beechey-Gradwell et al., 2020; Cooney et al., 2021). However, in our greenhouse experiment, this increase in leaf N concentration did not translate into clear evidence for substantially higher whole-plant N accumulation, particularly under drought conditions where reduced biomass appeared to offset concentration-level changes. Studies in high-lipid ryegrass further demonstrate that enhanced C allocation to vegetative lipids increases N uptake and redistributes N toward metabolic pathways that support growth and energy production (Beechey-Gradwell et al., 2018; Cooney et al., 2021). Similar mechanisms may contribute to the elevated leaf N concentrations we observed in oil-enhanced sorghum, where increased vegetative oil accumulation likely created greater demand for ATP, reductant, and C precursors (Roberts et al., 2025), which may have stimulated N assimilation and preferential leaf-level N allocation.

Despite the aboveground effects we observed, oil enhancement did not significantly alter belowground C allocation or rhizosphere C inputs under well-watered conditions (Fig. 8). Root biomass, root ^13^C enrichment, and soil ^13^C incorporation did not differ across genotypes, suggesting that changing the plant’s lipid sinks primarily influenced leaf-level nutrient and C dynamics rather than directing additional C belowground. Under drought, both sorghum genotypes experienced reductions in stomatal conductance, photosynthesis, leaf water status, and biomass, indicating that moisture-driven limitations on C assimilation largely overwhelmed genotypic differences in C allocation. Overall, our findings support **H1** by showing that engineered oil accumulation modified sorghum physiology, C allocation, and N partitioning under well-watered conditions, while drought exerted primary control over C partitioning and root–soil interactions. This moisture-dependent expression of engineered traits highlights the need to consider genotype by environment interactions when evaluating and selecting oil-enhanced sorghum for water-limited bioenergy systems.

### 4.2 Drought Effects on Sorghum Physiology and Carbon Allocation

Drought imposed a coordinated set of physiological limitations that reduced C assimilation and reshaped whole-plant resource partitioning across both sorghum genotypes, consistent with our second hypothesis (**H2)**. Lower soil moisture reduced stomatal conductance, steady-state photosynthesis, and leaf water content, which are well-established responses of C4 grasses experiencing moderate to severe water stress (Abreha et al., 2022). Drought also shortened stomatal length and modestly increased stomatal density, anatomical adjustments that restrict CO2 diffusion and enhance hydraulic safety (Driesen et al., 2023). These combined structural and functional constraints reduced shoot and root biomass and lowered plant height, and promoted higher intrinsic water-use efficiency, indicating that sorghum adopted a more conservative water use strategy as C assimilation became increasingly constrained by moisture limitation. We suspect that these drought-induced constraints minimized genotypic differences in aboveground traits, further emphasizing the overriding influence of soil moisture on physiological performance relative to variation in engineered sink strength.

In our study, drought significantly increased leaf, stem, and root N concentrations in sorghum (Fig. 4), a pattern that contrasts with many previous studies which have reported reduced plant N concentrations under water limitation (He and Dijkstra, 2014; Wang et al., 2014; Munene et al., 2025). Although declining soil moisture generally constrains N uptake (Bista et al., 2018; Plett et al., 2020), tissue N concentration reflects the balance between N acquisition and C assimilation. When drought suppresses photosynthesis and biomass accumulation more strongly than it limits N uptake, N can become concentrated within a smaller biomass pool, resulting in higher N per unit dry mass. Sorghum’s relatively high N-use efficiency under water stress (Wang et al., 2014) may further support maintenance of N acquisition or internal N recycling. Mycorrhizal symbioses may also enhance nutrient uptake during drought; a ^15^N tracer study showed that arbuscular mycorrhizal hyphae increased ^15^N uptake by 4–11.5 times under drought relative to well-watered sorghum (Munene et al., 2025), likely because hyphae span air-filled pores and access nutrient and water pools that become disconnected from roots as soils dry (Kakouridis et al., 2022). Furthermore, drought likely stimulated the synthesis of N-rich metabolic compounds, such as proline and other compatible solutes, which support osmotic adjustment and help stabilize metabolism under water stress (Hagedorn et al., 2016; Wang et al., 2022). The concurrent increase in NSC observed under drought aligns with evidence that carbohydrate accumulation contributes to osmotic buffering (Hagedorn et al., 2016). The strong positive correlation between root NSC and root N in our study indicates coordinated regulation of carbon storage and N-supported osmotic functions, an adaptive mechanism proposed to maintain metabolic stability when carbon supply is restricted but demand for N-containing osmolytes increases (Wang et al., 2022). Collectively, the simultaneous increases in root NSC, root N, and topsoil root ^13^C enrichment suggest that droughted sorghum preferentially allocated recently assimilated C into N-supported metabolic and storage pools that strengthen root resilience.

To our knowledge, no previous study has quantified depth-resolved ^13^C allocation simultaneously in plant roots and rhizosphere soil under drought, particularly in an annual bioenergy crop. In our study, we found that drought reduced the absolute amount of excess ^13^C retained in soil, but did not reduce the proportion of whole plant ^13^C assimilates allocated belowground, consistent with prior studies showing lower absolute but greater proportional soil C inputs under drought (Burri et al., 2013; Wang et al., 2021). More importantly, drought caused pronounced vertical changes in recent C allocation, supporting our third hypothesis (**H3)**. Under well-watered conditions, deeper roots had high ^13^C enrichment, indicating active downward transport of recent assimilates (Hoekstra et al., 2014; Maan et al., 2023). Under drought, this pattern reversed and shallow roots were more strongly enriched in ^13^C, while deeper roots showed markedly reduced enrichment. This pattern likely reflects reduced phloem transport efficiency combined with lower sink strength and limited metabolic activity in deeper, drier soil layers (Savage et al., 2016; Sevanto, 2018; Hartmann et al., 2020). Despite lower enrichment in deep roots, drought significantly increased ^13^C enrichment in deep rhizosphere soil. This divergence indicates a decoupling between C incorporation into deep root biomass and C deposition into deep rhizosphere soil, suggesting that drought enhances the transfer or retention of recent assimilates in deeper soil layers through rhizodeposition, fine-root turnover, or increased microbial use of mobile C compounds (Fuchslueger et al., 2022). The rhizodeposition index further supports this pattern, showing that drought lowered rhizodeposition in surface soil but increased the proportion of root-derived C retained in soil at depth (Fig. 8). These depth-specific shifts represent a previously underexamined pathway by which drought alters not only internal C distribution but also the routes through which recent assimilates enter the soil.

Overall, our study provides mechanistic support for **H2** and **H3**, showing that drought reorganizes both organ-level and vertical distributions of recent assimilated C in both sorghum genotypes, enhances root C storage, and alters rhizosphere C transfer across soil depths. These insights advance understanding of how water limitation shapes C flow from bioenergy crops into soil and improve predictions of sorghum performance and soil C dynamics under increasingly variable moisture conditions.

### 4.3. Integration of Genetic and Environmental Controls on Carbon Partitioning

Our results provide an integrated assessment of how engineered lipid sinks and water limitation together regulate C allocation within sorghum and across the plant–soil interface. Oil enhancement primarily influenced aboveground C and N dynamics under well-watered conditions, consistent with the expectation that strengthened foliar sinks reshape photosynthetic demand, N allocation, and short-term retention of recent assimilates. Importantly, these genotypic differences did not extend to belowground C fluxes, indicating that lipid-based metabolic modification largely restructures shoot-level processes without substantially altering whole-plant partitioning to roots or soil C inputs. Because we quantified retained ¹³C in plant and soil pools rather than respiratory ¹³C losses, these allocation estimates represent recovered recent assimilates rather than a complete whole-plant C budget. Nevertheless, drought fundamentally reorganized sorghum C allocation regardless of genotype, supporting the conclusion that moisture availability is the principal driver of both internal C distribution and the depth at which recent assimilates enter soil. The drought-induced decoupling of deep-root and deep-soil ^13^C enrichment reveals a novel, underappreciated mechanism through which moisture stress alters rhizosphere C transfer pathways. This result underscores the importance of depth-explicit measurements for predicting belowground C persistence in deep-rooted annual bioenergy crops. Overall, these findings demonstrate that engineered metabolic traits and drought interact in complex, moisture-dependent ways to control C use within sorghum and the depth at which recent assimilates enter soil. This study establishes a mechanistic foundation for predicting the biogeochemical consequences of growing oil-enhanced sorghum under reduced rainfall regimes and informs efforts to develop climate-resilient, high-value bioenergy cropping systems that maximize both biomass productivity and soil C accumulation.

## 5. Conclusions

This study demonstrates that moisture availability is an overriding determinant of whole-plant function and C allocation in bioenergy sorghum, whereas engineering the plant’s lipid sink primarily altered C allocation strategies, and only under well-watered conditions. Drought consistently suppressed photosynthetic rates, modified stomatal anatomy, reduced biomass accumulation, and shifted recent C investment toward root storage pools and rhizosphere soil, indicating that moisture limitation strongly constrains whole-plant function and belowground C fluxes. Furthermore, drought induced a critical reorganization of C flow across soil depths, decoupling C transport to deep roots from C deposition in the deep rhizosphere. This reveals a previously unrecognized mechanism through which moisture stress alters pathways for subsoil C transfer. In contrast, oil enhancement elevated leaf N concentration, strengthened incorporation of new C into foliage, and increased leaf biomass only when water was sufficient, demonstrating that engineered lipid accumulation influences source–sink relationships in a moisture-dependent manner. By combining ^13^CO₂ continuous labeling with coordinated measurements of plant physiology, root traits, and soil C pools, our experiment clarifies how an engineered trait and moisture conditions can interact to shape C flow in deep-rooted sorghum. These findings provide a foundation for anticipating bioenergy sorghum productivity and soil C outcomes across increasingly variable moisture conditions, underscoring that moisture-driven constraints are the key bottleneck limiting the translation of metabolic innovations into robust, ecosystem-level benefits.

## Author Contributions

**Y. Liu**: Conceptualization, Methodology, Investigation, Data curation, Formal analysis, Visualization, Writing original draft. **K. Estera-Molina**: Investigation, Methodology, Data curation, Writing review and editing. **M. Kan**: Investigation, Methodology, Data curation, Writing review and editing. **J. Masterson**: Investigation, Writing review and editing. **C. Ramon**: Investigation, Writing review and editing. **A. Curtright**: Methodology, Writing review and editing. **R. Patra:** Investigation, Writing review and editing. **L. P. de Oliveira**: Investigation, Methodology, Formal analysis, Writing review and editing. **K. Park**: Investigation, Methodology, Writing review and editing. **E. Cahoon**: Resources, Writing review and editing. **W. Yang**: Conceptualization, Supervision, Writing review and editing. **J. Pett-Ridge**: Conceptualization, Supervision, Funding acquisition, Writing review and editing.

## Acknowledgements

We thank Aaron Chew, Manisha Dolui, Allegra Mayer, Joshua White, Zhen Li, Mike Allen, Kristina Rolison, Rhona Stuart, Steve Blazewicz, Noah Sokol, Kenzo Esquivel, Paul Zander, Shannon Brown, Erin Nuccio, Linnea Hernandez, Jessica Wollard, Di Liang, Changfeng Zhang, Hansen Qian, Stewart Anthony John, Anh Trinh, Cinthia Silveira, Keith Morrison, Mary Firestone, Hannah Trinh, Stella Ho, Zoe Woodhouse, Irene Yzabelle Fernandez, Melanie Rodriguez Fuentes, Emily Catherine Savrnoch, Tamara Miller, Leylen Miloslavich, Maggie Yuan, Elly Stone, Tracy Johnson for assistance with soil collection, greenhouse harvests and lab analyses; Christina Wistrom at the Oxford Tract Greenhouse at UC Berkeley where the experiment was conducted, and John Bailey, Troy McWilliams, and Greg Solberg at the Hopland Research and Extension Center, where soil was collected. We thank Brad Erkkila at the Yale Analytical and Stable Isotope Laboratory for assistance with stable isotope analyses. Tracy Johnson (University of Illinois Urbana-Champaign) coordinated the transfer of the sorghum seeds and Michelle Zambrano (Lisa Ainsworth’s lab, University of Illinois Urbana-Champaign) performed root non-structural carbohydrate analyses. We gratefully acknowledge Chunhwa Jang and DoKyoung Lee (University of Illinois Urbana-Champaign) for providing the sorghum seed materials used in this study. We thank Tara Nazarenus and Mengyuan Wang (Edgar Cahoon’s group) at the University of Nebraska-Lincoln for coordinating USDA requirements for receiving samples for root lipid analysis. This work was funded by the DOE Center for Advanced Bioenergy and Bioproducts Innovation (U.S. Department of Energy, Office of Science, Biological and Environmental Research Program under Award Number DE-SC0018420, and LLNL subaward SCW1798). Work conducted at LLNL was conducted under the auspices of the U.S. Department of Energy under Contract DE-AC52-07NA27344. Any opinions, findings, and conclusions or recommendations expressed in this publication are those of the author(s) and do not necessarily reflect the views of the U.S. Department of Energy.

## Conflicts of Interest

The authors declare no conflicts of interest.

## Notes

### Competing Interest Statement

The authors have declared no competing interest.

