## Supplemental Table and Figures for "Drought Dominates Engineered Lipid Sink Effects on Sorghum Physiology and Carbon Allocation"

| | Water condition | Sorghum type | Water condition x Sorghum type |
|---|---|---|---|
| Plant height | 31.65*** | 0.53 | 0.67 |
| Shoot biomass | 171.91*** | 0.65 | 0.89 |
| Stem biomass | 171.91*** | 0.65 | 0.89 |
| Leaf biomass | 90.43*** | **6.78*** | 5.06* |
| Root biomass | 171.91*** | 0.21 | 0.018 |
| Root:Shoot | 10.70** | 0.044 | 0.14 |
| GasEx | 92.09*** | 1.07 | 2.86 |
| iWUE | 3.36† | 0.015 | 2.17 |
| gsw | 701.8*** | 1.84 | 2.84 |
| Adaxial_SD | 2.39 | **3.03†** | 0.91 |
| Abaxial_SD | 0.99 | **3.02†** | 1.23 |
| Average_SD | 1.76 | **3.46†** | 1.22 |
| Adaxial_SL | 5.79* | **3.36†** | 0.023 |
| Abaxial_SL | 4.45* | 0.52 | 0.95 |
| Average_SL | 5.97* | 1.91 | 0.37 |
| Leaf_RWC | 5.69* | 0.51 | 0.001 |
| SLA | 4.37* | 0.001 | 0.37 |
| Leaf C | 0.037 | **4.17†** | 0.39 |
| Leaf N | 9.24** | **4.42*** | 0.91 |
| Leaf C: N | 10.77** | **5.89*** | 1.80 |
| Stem C | 0.97 | **3.14†** | 0.03 |
| Stem N | 14.64*** | 0.25 | 0.60 |
| Stem C: N | 10.78** | 0.65 | 0.52 |
| Root C | 1.75 | 0.96 | 2.18 |
| Root N | 123.39*** | 0.112 | 0 |
| Root C: N | 109.29*** | 0.091 | 0.73 |
| Leaf_enrichment | 4.83* | 0.004 | 1.61 |
| Stem_enrichment | 2.15 | 0.12 | 0.2 |
| Root:A_enrichment | 21.17*** | 0.034 | 0.05 |
| Root:B_enrichment | 0.164 | 0.063 | 0.016 |
| Root:C_enrichment | 7.88* | 0.53 | 0.13 |
| Soil:A_enrichment | 1.76 | 0.27 | 0.72 |
| Soil:B_enrichment | 1.91 | 0.93 | 0.14 |
| Soil:C_enrichment | 4.58* | 0.13 | 0.044 |

**Table S1** Summary of two-way ANOVA results (*F*-values and significance levels) for sorghum physiological, morphological, and biogeochemical parameters. Symbols: *** $p < 0.001$; ** $p < 0.01$; * $p < 0.05$; † $p < 0.1$; ns = not significant.

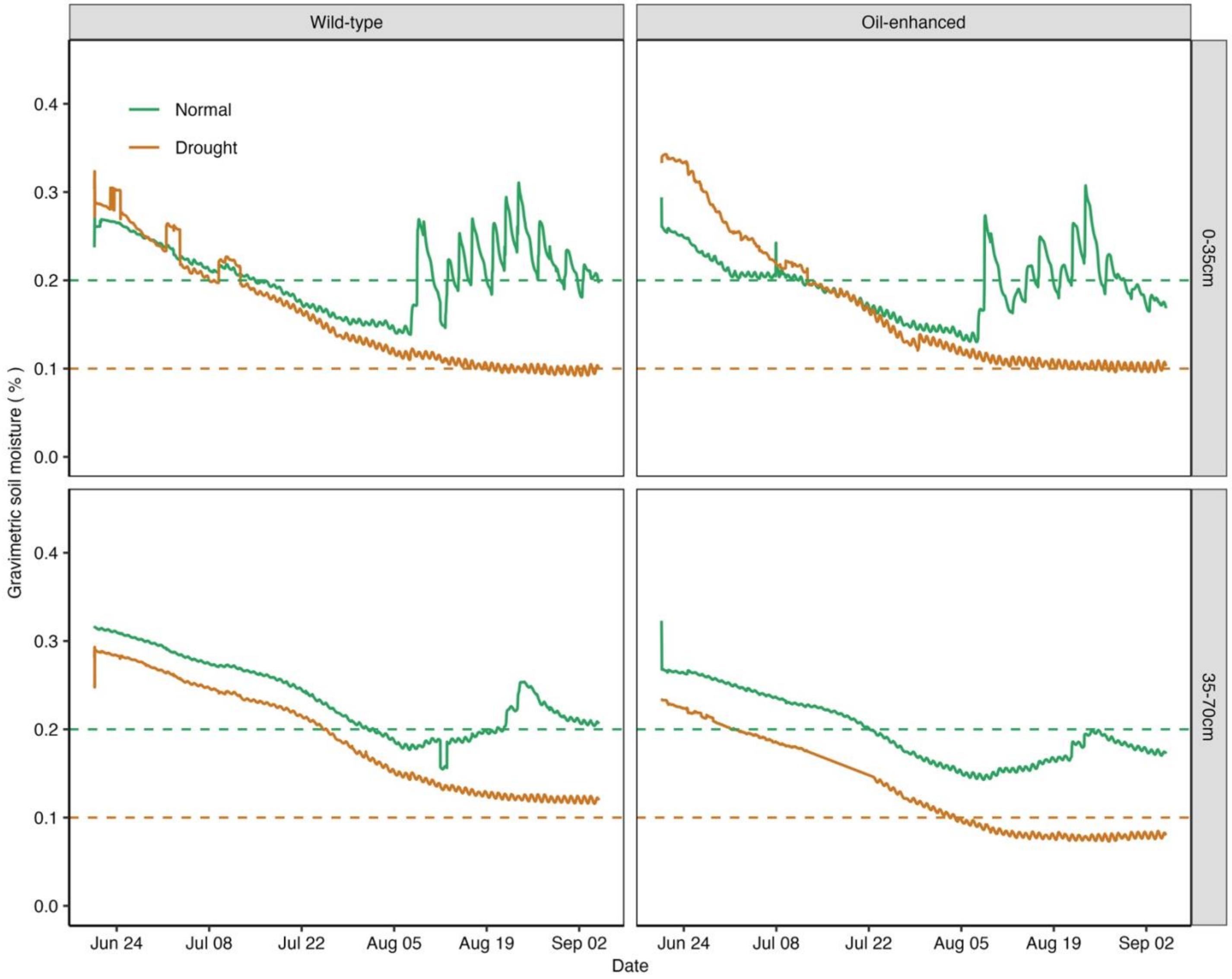


**Fig. S1 Temporal dynamics of soil gravimetric water content (GWC).** Soil GWC at two depths (0–35 cm, top panels; 35–70 cm, bottom panels) for wild-type (left) and oil-enhanced (right) sorghum grown under normal (green) and drought (brown) moisture treatments. Dashed horizontal lines represent average GWC levels maintained for each treatment.

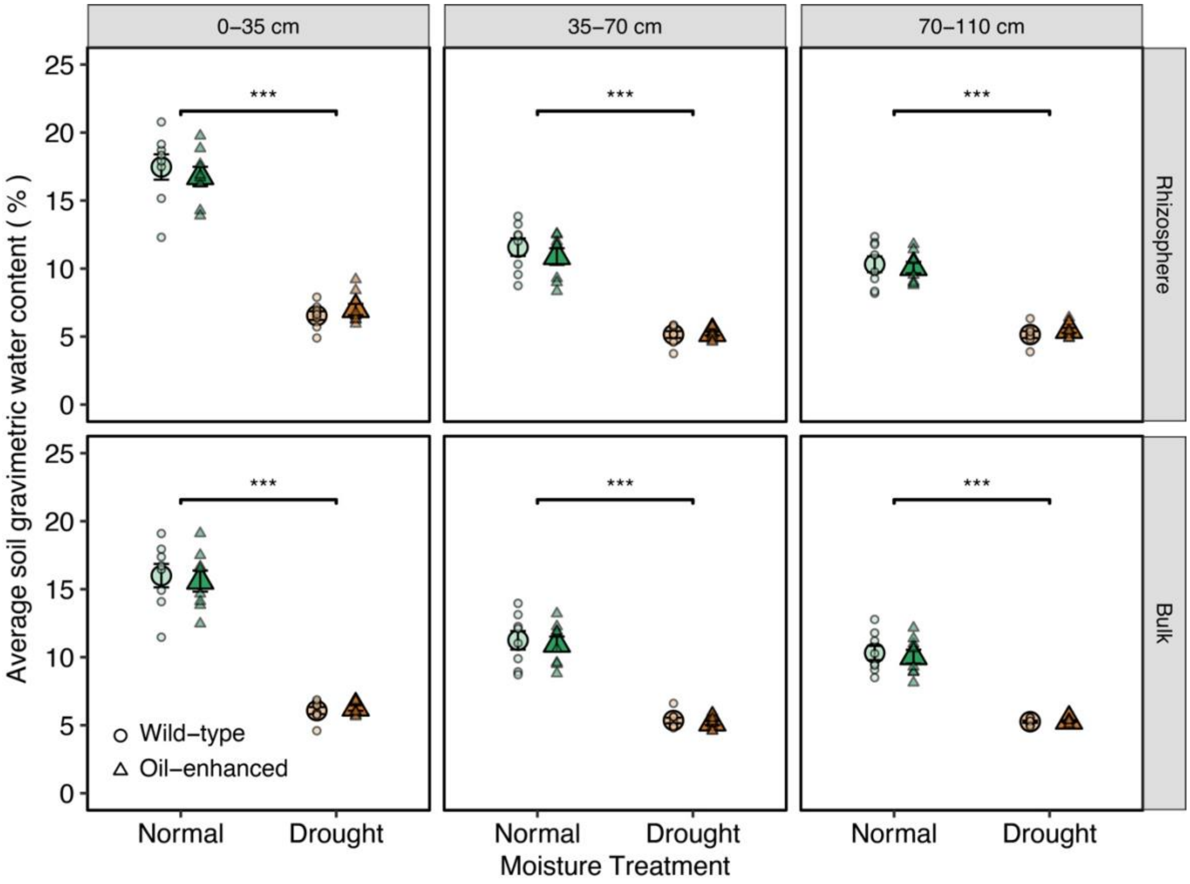


**Fig S2. Average soil gravimetric water content (GWC) across depths under normal and drought conditions.** Average soil GWC (%) measured in the rhizosphere (top panels) and bulk soil (bottom panels) at three depths (0–35 cm, 35–70 cm, and 70–110 cm) for wild-type (circles) and oil-enhanced (triangles) sorghum under normal (green) and drought (brown) moisture treatments. Bars represent means ± SE (n = 8). Asterisks indicate significant differences between moisture treatments (*** $p < 0.001$).

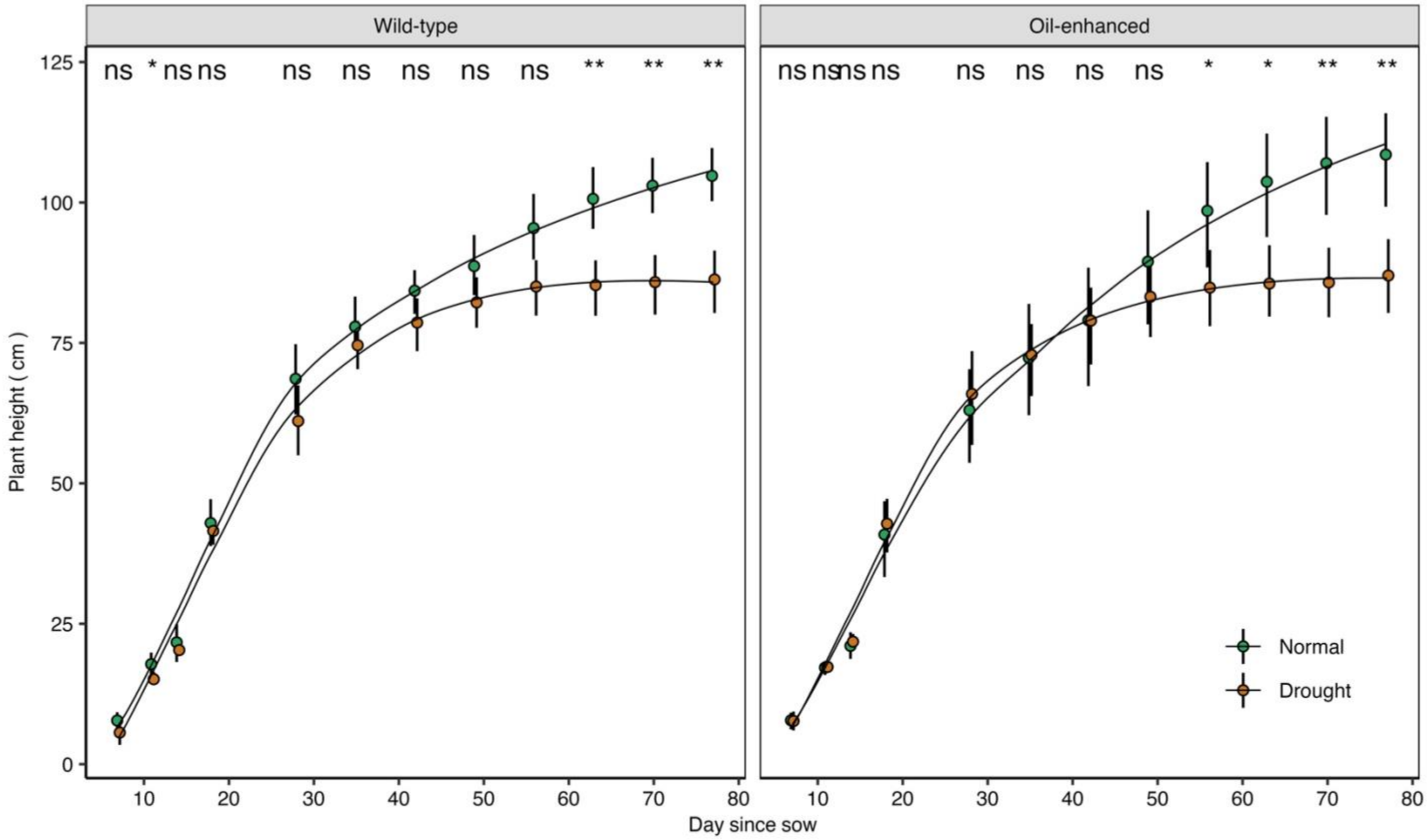


**Fig. S3 Growth dynamics of sorghum plant height under normal and drought conditions.** Time course of plant height (cm) for wild-type (left) and oil-enhanced (right) sorghum grown under normal (green) and drought (brown) moisture treatments. Points represent means ± SE (n = 8). Asterisks indicate significant differences between moisture treatments at each time point (* $p < 0.05$, ** $p < 0.01$, ns = not significant).

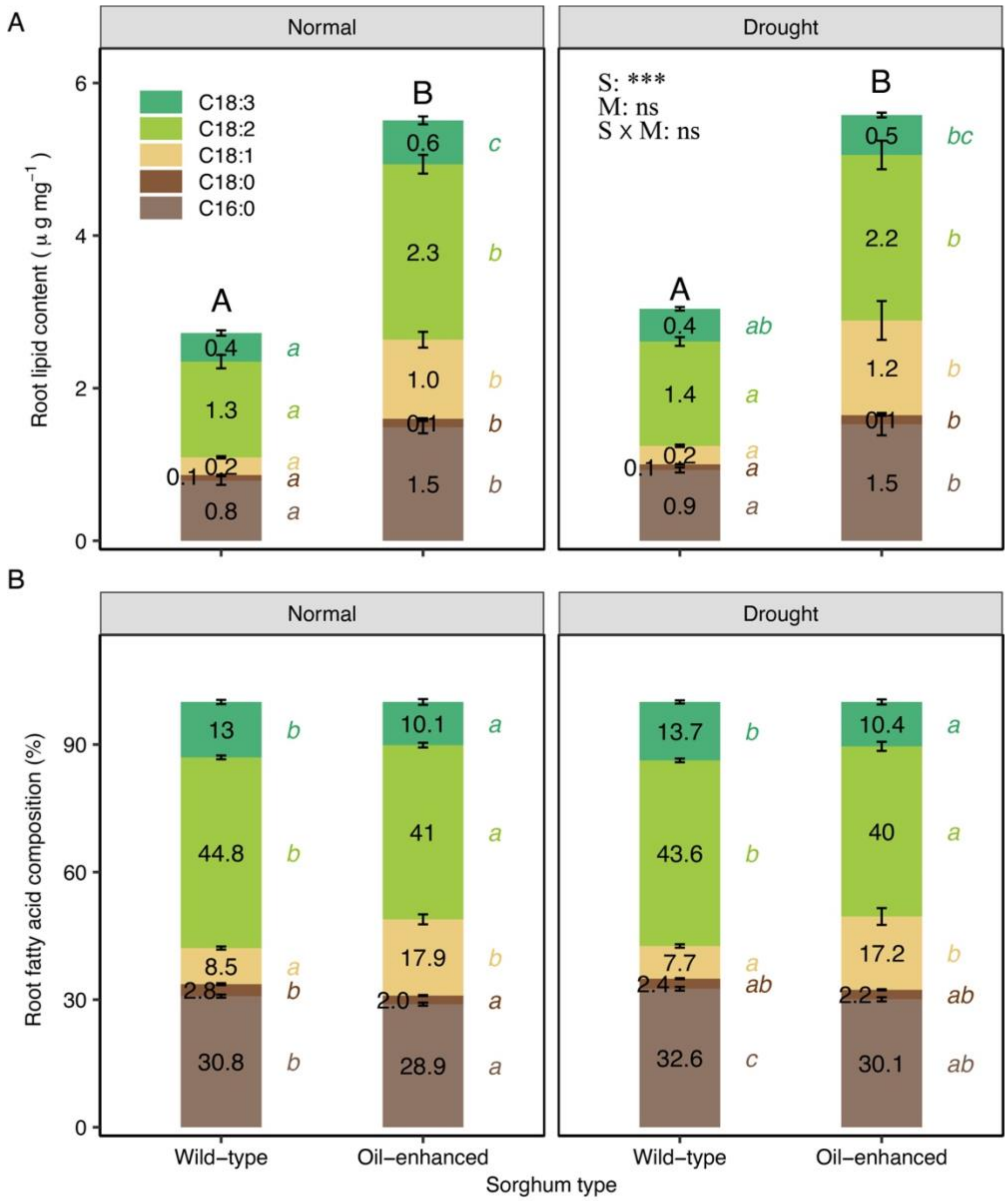


**Fig. S4 Effects of oil enhancement and drought on root lipid content (A) and fatty acid composition (B).** Total lipid content (µg mg$^{-1}$ dry mass) and proportional lipid composition (%) in wild-type and oil-enhanced sorghum grown under normal and drought moisture conditions across three horizons. Bars represent means ± SE (n = 24). Different uppercase letters above bars indicate significant differences in total lipid amount between sorghum types and moisture treatments, while lowercase letters indicate significant differences in individual fatty acid fractions based on two-way ANOVA followed by post hoc tests ($p < 0.05$).

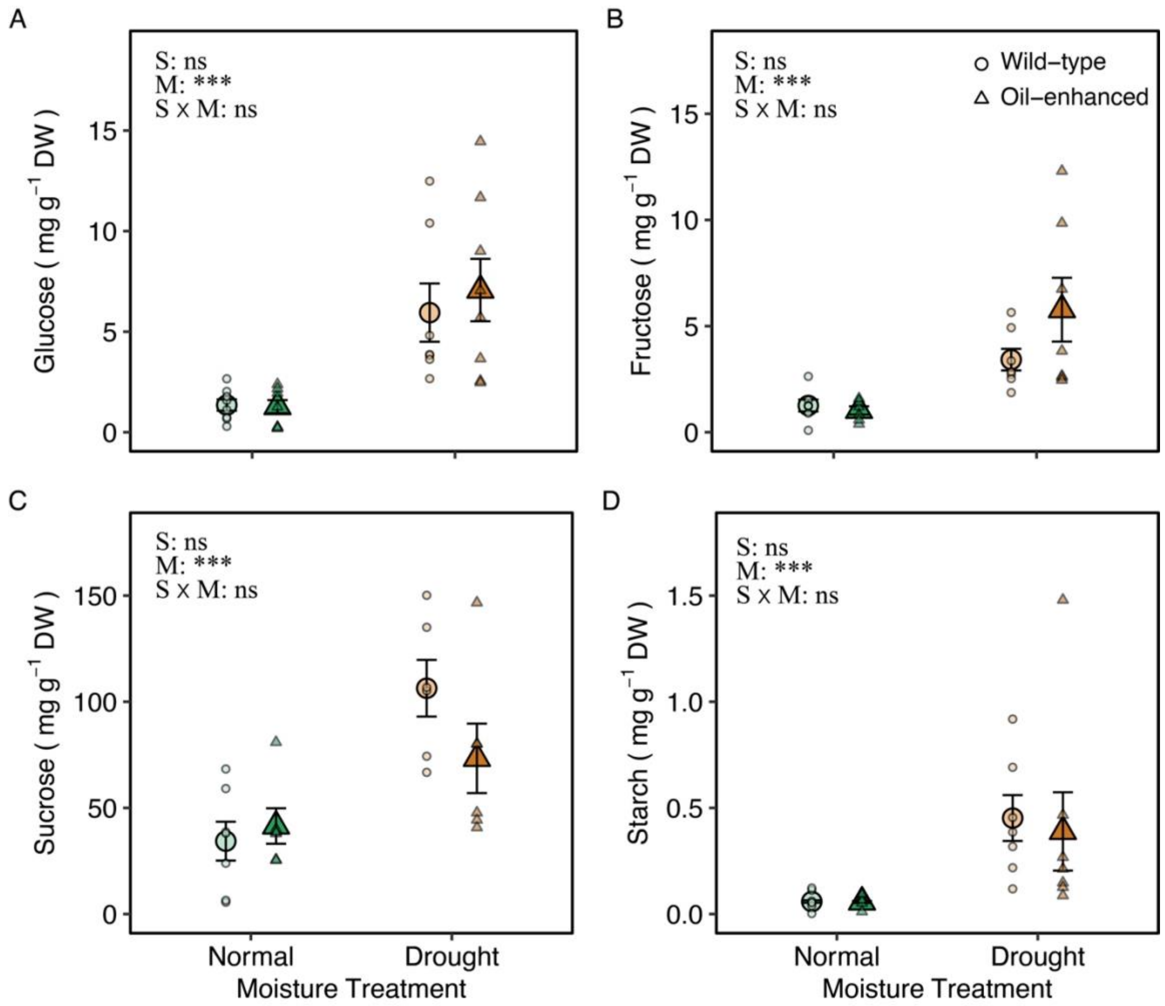


**Fig. S5 Effects of drought and oil enhancement on root non-structural carbohydrate pools in sorghum.** Root concentrations of (A) glucose, (B) fructose, (C) sucrose, and (D) starch are shown for wild-type and oil-enhanced sorghum grown under normal and drought moisture treatments. Points represent individual biological replicates; enlarged symbols denote mean ± SE. Circles and triangles indicate wild-type and oil-enhanced sorghum, respectively. Green and brown colors denote normal and drought treatments. Two-way ANOVA results are reported in each panel for effects of sorghum type (S), moisture treatment (M), and their interaction (S × M). Symbols: * $p < 0.05$; ** $p < 0.01$; *** $p < 0.001$; ns, not significant.

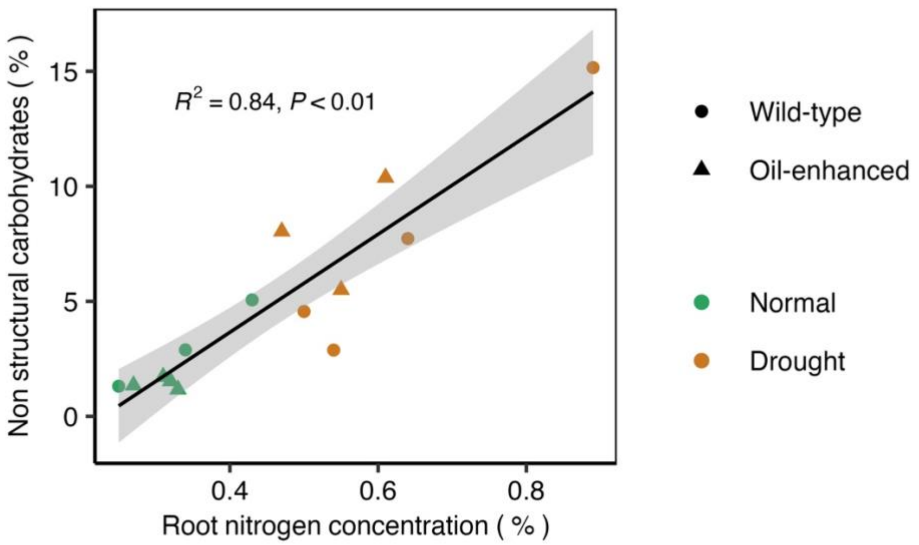


**Fig. S6 Relationship between root nitrogen concentration and non-structural carbohydrates in sorghum roots.** The solid line represents the linear regression fit, with the shaded area indicating the 95% confidence interval.

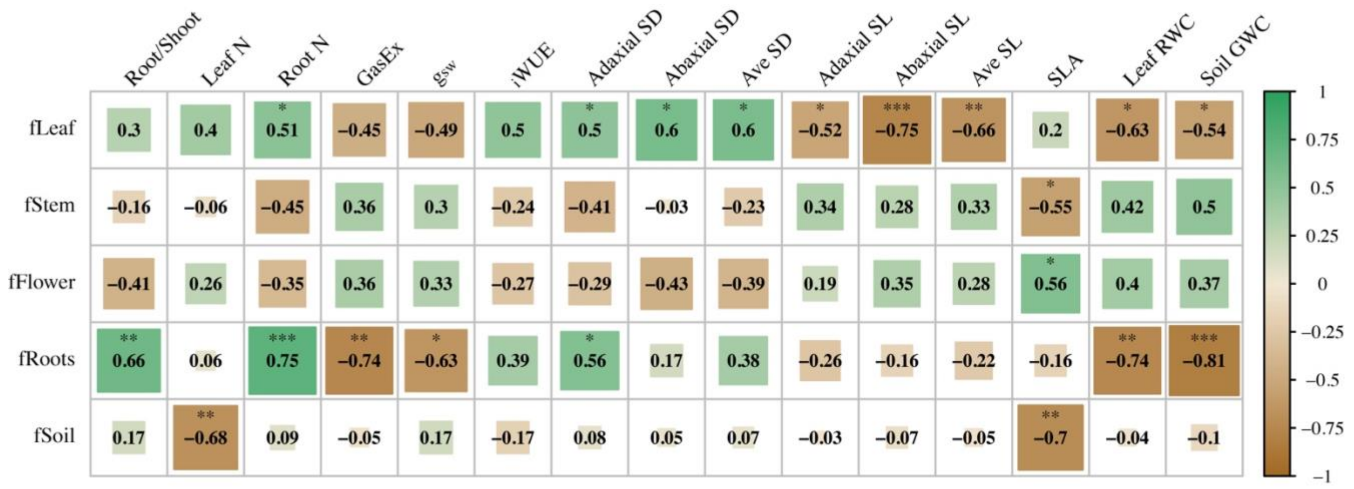


**Fig. S7 Correlation matrix between the fraction of new $^{13}C$ in plant tissues and soil with plant physiological and environmental traits.** The matrix shows Pearson correlation coefficients between the fraction of new $^{13}C$ (fLeaf, fStem, fFlower, fRoots, fSoil) and plant traits, including root-to-shoot ratio, leaf and root N concentration, steady state photosynthesis rate (GasEx), stomatal conductance ($g_{sw}$), water-use efficiency (iWUE), stomatal density (SD), stomatal length (SL), specific leaf area (SLA), leaf relative water content (RWC), and soil gravimetric water content (GWC). Color shading represents correlation strength and direction (green = positive, brown = negative), with significance levels indicated as * $p < 0.05$, ** $p < 0.01$, *** $p < 0.001$.
